# Parent-of-origin phasing of somatic mutations shows equal mutation burden between parental genomes in human cancers

**DOI:** 10.64898/2026.09.02.748867

**Authors:** Maxime Lefèbvre, Arnaud Cléris, Mathieu Parmentier, Peter Van Loo, Vincent Detours, Maxime Tarabichi

## Abstract

Somatic mutations accumulate independently in the two parental genome copies of our cells throughout life and shape cancer evolution. Although local mutation rates are influenced by allele-specific features such as DNA sequence, epigenetic marks, and chromatin structure, whether these translate into genome-wide differences in mutation accrual between the two parental copies is unknown. Cancer genomics analyses, including copy-number gain timing and molecular archaeology, assume that mutations accrue symmetrically on the two homologous parental genomes, yet this assumption has never been tested. Here we present **PhaSoMix**, a framework exploiting the genetic differentiation between parental haplotypes in admixed cancer patients to assign somatic mutations to their parent of origin without parent or parent-surrogate sequencing. Applying it with explicit modeling and propagation of phasing and ancestry-inference uncertainty across 21 tumor whole genomes from the Pan-Cancer Analysis of Whole Genomes cohort, we find mutation burdens highly symmetric between maternal and paternal genomes, across cancer types, genomic annotations, clonal timing categories, and mutational processes including clock-like CpG sites, bounding any asymmetry to within 4-5%. Simulations show that violations would substantially bias gain-timing estimates in late evolutionary windows. This provides the first quantification of parental mutation-burden symmetry *in vivo*, validating a key assumption of cancer evolutionary analyses.

## Introduction

Over the course of an individual’s lifetime, normal human cells accumulate somatic mutations, with documented rates of single nucleotide variants (SNVs) across human tissues^1^. The majority of these mutations are passengers and would not have large effects, while some driver mutations can alter key cellular functions^2^. Early somatic mutations can cause developmental disorders, whereas the progressive accumulation of mutations throughout life can lead to cancer and contribute to aging and age-related diseases^3^. Mutation rates vary among somatic cell types within the same individual, influenced by cell division rates and the inherent characteristics of each cell type^1^. They are modulated by environmental exposures, genetic predispositions, cellular stresses, and defects in DNA repair mechanisms. Within the same nucleus, the susceptibility to somatic mutations can also vary spatially along the genome.

In the nucleus of most healthy diploid somatic cells, two homologous copies of the human genome coevolve, one inherited from each parent. While somatic mutations occur independently on each of these two haploid genomes, it is believed that the two homologous parental genomes accumulate mutations at the same rates. This assumption is implicit in almost all cancer genomics analyses, thus violations could disrupt many genomewide estimates. For example, in molecular archaeology and gain timing analyses^4,5^, copy-number gains are leveraged as providential events for tumor evolution analyses, as they take a molecular snapshot of the DNA sequence at a given point in time. This snapshot can be leveraged to time the gains and the underlying mutations relative to that gain event^4,5^. One of the implicit assumptions in these analyses is that all homologous copies, including the two parental alleles, have the same somatic mutation rates over time locally and globally across chromosomes, thus the accrual should be equal. This assumption and the effects of its violation in cancer genomics analyses have never been tested.

Allelic differences include variations in DNA sequence, methylation, histone modifications, gene expression, and chromatin structure^6–8^, and each of these allele-specific factors has been shown to contribute to variations in local mutation rates^2,9–12^. While these effects may either decrease or increase mutation rates, they are expected to act without directional bias between maternal and paternal genomes. However, some less appreciated genome-wide parent-of-origin effects have been reported, manifesting transiently during embryogenesis, such as in replication timing and chromatin compartmentalization in mice^13^, or even persisting across generations, as seen in methylation signatures outside imprinted regions that reflect parental environmental exposures, including obesity, cold winters, or starvation^14^. Such factors, as well as others yet to be identified, could in principle alter mutation rates asymmetrically between the two parental genomes, for example by impacting the number of (C>T)pGs linked to methylation leakage during epigenetic resets on one parental copy. These genomewide parent-of-origin rates, however, have yet to be quantified.

Here, we aimed to explore the differences in somatic mutation accrual between the two parental copies of the human genome having coevolved in the same nuclei over time *in vivo*. Somatic mutations called from whole-genome sequencing in cancer tissues provide a high mutational load we can leverage to quantify this effect. However, while parent-of-origin whole-genome phasing is best addressed through approaches such as family trio sequencing, strand-seq and long-read sequencing^15^, or through parent-surrogate phasing^16^, existing cancer datasets lack the necessary data.

To address this, we present PhaSoMix, a methodology based on publicly available data that assigns somatic mutations in admixed cancer genomes to their parental haplotype of origin. Where haplotypes carry distinct local ancestries, ancestry-discriminant genomic segments serve as a proxy for parental origin, enabling quantification of mutational differences between parental chromosome sets. We apply PhaSoMix to the Pan-Cancer Analysis of Whole Genomes (PCAWG) cohort and propagate errors at each algorithmic step to bound the detectable difference in mutation accrual between the two parental genomes.

## Results

### PhaSoMix: a generic approach leveraging local ancestry to phase somatic mutations to parental genomes in admixed cancer patients

In the absence of parental genomes for genome-wide phasing in cancer data, we reasoned that recently admixed patients could be leveraged, as the genetic distance between their two parental genomes is maximized and ancestry assignment of local haplotypes can act as a stand-in for parent-of-origin assignment. Leveraging SNP-in-read phasing will then phase somatic mutations to local haplotypes, and thereby can assign ancestry, i.e. parent of origin.

We thus sought to develop PhaSoMix (**Methods**), a framework performing the following four steps: 1. Run local ancestry inference genome-wide across the PCAWG cancer cohort to identify exploitable patients, defined as those whose two parental haplotypes carry distinct ancestral origins across at least a fraction of the genome, creating ancestry-discriminant segments that serve as a proxy for parental identity (**Figure 1A**); 2. Empirically phase somatic mutations to nearby SNPs/haplotypes using raw sequencing data (**Figure 1B**); 3. Identify maternal and paternal ancestries from mitochondrial (mt) DNA and chromosomes X/Y SNPs (**Figure 1C**); 4. Aggregate phased somatic mutations by inferred parental origin to compute genome-wide parental mutation burdens, corrected for allele-specific copy number alterations (**Figure 1D**).

**Figure 1.**
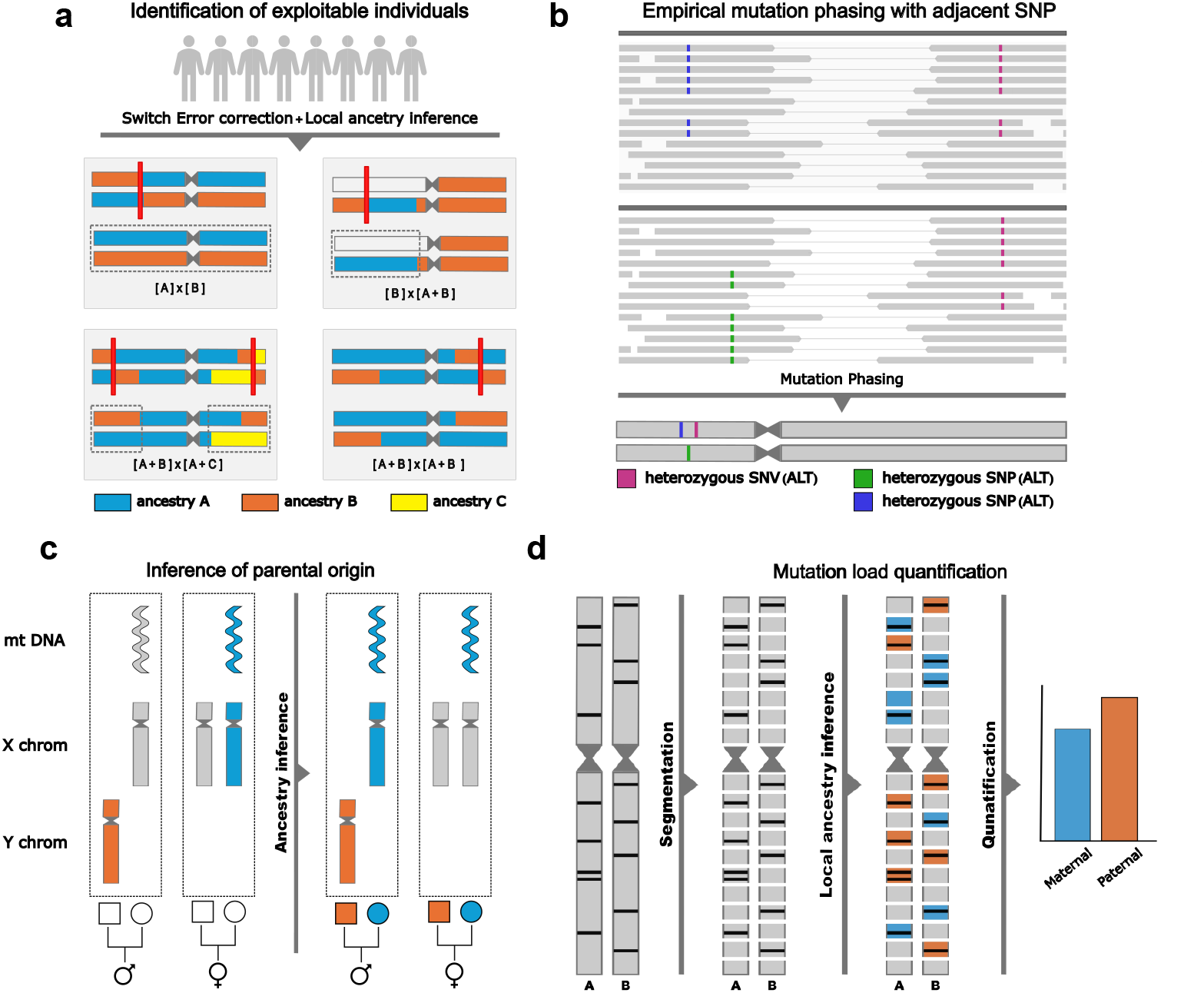
Overview of the PhaSoMlix framework for quantifying parental-specific somatic mutation rates. **(A)** Identification of informative individuals in the PCAWG cohort using haplotype-level local ancestry inference (Gnomix). Each haplotype is represented as a sequence of ancestry blocks (orange, blue, yellow). Switch errors (SE, red lines) are corrected in regions of allelic imbalance. Individuals are classified into four cases: a strict F1 individual with two fully mono-ancestral haplotypes (A x B); an individual with one mono-ancestral and one admixed haplotype (A x A+B); an individual where both haplotypes are admixed but derive from distinct ancestries (A+B x A+C); and a non-informative case where both haplotypes share the same ancestries (A+B x A+B). Dashed grey boxes highlight exploitable regions (genomic segments where the two haplotypes carry distinct, unambiguous ancestries) which serve as the analytical units for downstream comparisons. **(B)** Empirical phasing of somatic mutations onto parental haplotypes by linking them to nearby heterozygous SNPs using raw paired-end sequencing reads (grey blocks). **(C)** Inference of the maternal and paternal origin of each haplotype from mitochondrial DNA and sex chromosome (X/Y) variants, using an XGBoost classifier. **(D)** Computation of genome-wide parental-specific mutation burdens by aggregating phased somatic mutations according to their inferred parental origin, with counts corrected for allele-specific copy number alterations. Black horizontal ticks represent individual somatic mutations.

Parent-of-origin attribution is only possible where a given ancestry uniquely identifies one parent, which depends on each patient’s admixture profile. We distinguish four informative cases (**Figure 1A**). In the first, both parents are mono-ancestral with different ancestries (e.g. [A] × [B]): every genomic segment maps unambiguously to one parent and the entire genome is exploitable. In the second, one parent is mono-ancestral and the other parent admixed (e.g. [A] × [A+B]): only segments carrying an ancestry unique to the admixed parent permit attribution. In the third, both parents are admixed with partially overlapping ancestries (e.g. [A-B] × [B-C]): ancestries unique to each parent allow attribution, while the shared ancestry is excluded. In cases 2 and 3, only ancestry-discriminant regions are used.

At each step of this pipeline, however, errors accumulate, which add noise to our estimates. We therefore designed experiments to measure the noise for each of those steps, which allowed us to propagate errors.

### Measuring and propagating errors to obtain upper-bound measurements on the difference in mutation rates between the parental genomes

To measure and be able to propagate errors in our pipeline, we next designed simulation experiments leveraging self-reported and inferred ancestries from the 1,000 genomes project (1kGP) and derived ground truth phasing from copy-number imbalances in cancer genomes (**Figure 2**; **Methods**).

**Figure 2.**
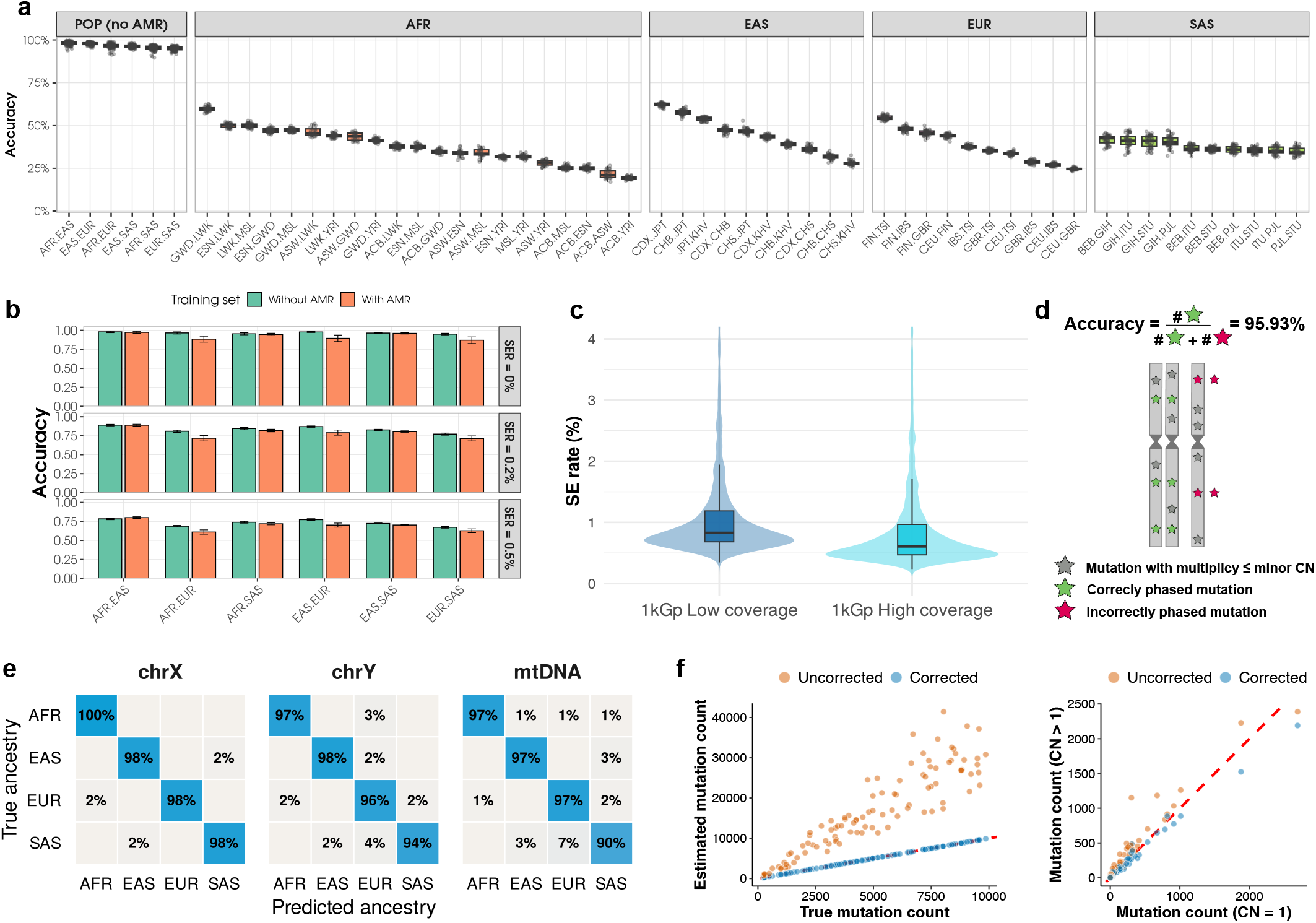
Evaluating stepwise accuracy of PhaSoMix to quantify propagated error. **(A)** Chromosome-pair accuracy of local ancestry inference (Gnomix) at SER = 0%. Each panel corresponds to a super­population (AFR, EAS, EUR, SAS) or the full population set excluding AMR (POP no AMR). Within each panel, boxplots show the distribution of weighted mean accuracy across chromosomes 1-22 for each ancestry pair combination, ordered by decreasing median accuracy. Individual points represent replicate simulations. **(B)** Accuracy of local ancestry inference (Gnomix) across 1000 Genomes Project super-populations (AFR, EAS, EUR, SAS), evaluated with and without the AMR population in the training set, and under increasing switch error rates. Each bar represents mean accuracy per super-population, with and without AMR included in the training set (green and orange, respectively). Three switch error rate (SER) conditions are shown: SER = 0% (top), SER = 0.2% (middle), and SER = 0.5% (bottom). **(C)** Switch error rate (SER) for PCAWG samples phased with Beagle 5.4 using two different reference panels: low-coverage and high coverage 1kGp reference panels. **(D)** Accuracy of SNV phasing in allelic-imbalance regions, measured as the proportion of SNVs whose inferred multiplicity does not exceed the copy number of the phased allele. **(E)** Confusion matrices for XGBoost-based parental origin classification across chrX, chrY, and mtDNA. Each cell shows the proportion of samples with a given true ancestry (rows) assigned to each predicted ancestry (columns), normalized by true class. Super-populations follow the 1000 Genomes Project classification (AFR, EAS, EUR, SAS). **(F)** Effect of copy-number correction on local mutation counts per Mb. Left panel: evaluation using simulated data with controlled mutation rates and copy-number alterations. Right panel: assessment in PCAWG segments with copy-number gains and losses.

First, we evaluated the accuracy of local ancestry inference using Gnomix ^17^ (**Figure 2; Extended Data Figure S1)**. Accuracy was assessed through simulations of admixed genomes generated from 1kGP and PCAWG data for which the assignment of haplotype to ancestry is known by design, with increasing switch-error rates (SER) and, covering all pairwise combinations both at the super-population level: African (AFR), Admixed American (AMR), East Asian (EAS), European (EUR), and South Asian (SAS); and at the sub-population level within each super-population (**Methods**). Sub-population resolution proved less reliable, with accuracies too low to support downstream analyses (**Figure 2A**). We therefore restricted local ancestry inference to the super-population level, with mean accuracy ranging from 95% to 98% across populations (**Figure 2A**). However, we observed that including AMR substantially degraded accuracy, due to lower representation of AMR in the 1kGP dataset, and particularly for combinations involving EUR (**Figure 2B**), consistent with the fact that AMR samples are themselves historically admixed and harbour a large EUR component (**Extended Data Figure S2 A-B**). We thus excluded AMR from the reference set and retained the four remaining super-populations (AFR, EAS, EUR, SAS) for all downstream analyses.

Because somatic phasing errors are expected to degrade local ancestry inference, we further assessed the robustness of our predictions to phasing noise by simulating increasing switch error rates. In simulations, accuracy dropped sharply as the simulated SER increased, most notably for the AFR–EUR combination, where it fell from 96.54% at SER = 0%, to 80.7% at SER = 0.2%, and only 68.65% at SER = 0.5% (**Figure 2B**). This steep decay underscores the need for high-quality, accurately phased haplotypes. In admixed genomes, a single switch error creates a segment containing alleles from both parental ancestries, directly confounding local ancestry inference. To achieve the best possible phasing, we updated the reference 1kGP genotypes with the more recent high-depth release (**Methods**). This improvement reduced switch error rates by 26.98%, reaching a median SER of 0.6% (**Figure 2C**).

While this represents a substantial improvement, residual switch errors still remain in genomic regions where phasing cannot be independently validated. We next focused our benchmark on allelic-imbalance regions in real data, where copy-number information allows residual switch errors to be corrected to near-perfect phasing, providing the high-confidence haplotype assignments required for reliable local ancestry inference and parent-of-origin attribution (**Extended Data Figure S3**; **Methods**). First, phasing accuracy of SNVs to nearby SNPs in the informative PCAWG patients was evaluated in regions of allelic imbalances, where phasing error rates can be calculated as the fraction of SNVs phased to the minor allele with an inferred multiplicity exceeding the number of copies of the minor allele^18^. Using this metric, we observed a high overall agreement of 95.93%, indicating that the vast majority of SNVs were correctly assigned to their respective haplotypes by our pipeline (**Figure 2D; Methods**).

Second, given the low accuracy at high SER in simulations, we asked how much residual phasing noise actually affects final SNV parent-of-origin assignments in real data. We first compared assignments obtained with and without switch-error correction in imbalanced regions, and found them highly concordant (median 92%) despite the reduced accuracy expected at this switch-error rate in simulations. To confirm this directly, we introduced simulated switch errors into the corrected imbalance regions and recomputed assignments, again observing high but lower concordance (median 85%; **Extended Data Figure S4**). The lower value likely reflects differences in how simulated and real switch errors are distributed across the genome, indicating that SNV parent-of-origin assignment is more robust to residual phasing noise in real data than in our simulations.

Parental haplotypes in admixed individuals were assigned from mtDNA SNPs, with additional resolution from chromosome X and chromosome Y in males using XGBoost models trained on 1kGP data (**Methods**). Across the four super-populations (AFR, EAS, EUR, SAS), accuracy was uniformly high on chrX (98–100%) and chrY (94–98%) (**Figure 2E)**. On mtDNA, AFR, EAS and EUR reached 97% accuracy, while SAS showed a modestly lower rate of 90%, with the remaining 10% misassigned primarily to EUR (7%) and EAS (3%). Diagonal dominance across all three confusion matrices (**Figure 2E**) confirms that chrX, chrY and mtDNA each carry sufficient signal to robustly assign paternal and maternal ancestries. Consistent with the local ancestry inference on autosomes, including the AMR population in the training set decreased prediction accuracy from chrX, chrY and mtDNA (**Extended Data Figure S5**).

Copy-number gains and losses occur throughout tumor evolution ^5^ and alter the cumulative mutation rate per ancestral copy. We applied a correction formula to adjust mutation counts for the number of copies. In simulations, this correction recovered the true mutation rate with no bias, regardless of copy number (**Figure 2F; Methods**). In real PCAWG data, uncorrected counts increased with copy number, as expected, though less steeply than in simulations, potentially because low-multiplicity mutations are less likely detected by mutation-calling pipelines at higher copy number (**Supplementary Table 2)**. After correction, we observed a modest residual overcorrection (1–7%) at copy number states 2, 3 or 4, the most represented states in the dataset. Given this residual overcorrection in imbalanced regions, mutation rates were quantified separately in balanced and imbalanced regions to test our original hypothesis.

By quantifying the accuracy at each step of our pipeline, we can propagate the cumulative error on our mutation burden estimates, affected by the accuracy of the assignment of SNVs to their parental haplotype of origin. This calculation therefore incorporated the SNV phasing accuracy, and the calculated accuracy of local ancestry inference (**Methods**).

### PhaSoMix identifies 21 informative admixed genomes in the 2,645-patient PCAWG cohort

We focused our analysis on the Pan-Cancer Analysis of Whole Genomes (PCAWG) dataset, which provides high-quality consensus somatic mutation calls. We phased all 2,645 genomes at 1kGP SNP positions using Beagle 5.4 with a high-coverage 1kGP reference panel, and corrected residual switch errors within allelic-imbalance regions (**Methods**). We then ran local ancestry inference genome-wide with Gnomix ^19^ across the full cohort and used the per-haplotype ancestry profiles to identify informative patients, defined as those in whom at least one ancestry uniquely tags a single parent (**Methods**). Patients with internally inconsistent or ambiguous ancestry assignments were excluded.

This yielded 21 informative individuals among the 2,645 PCAWG patients (**Figure 3A; Supplementary Figures S1-20**). One was an F1-admixed individual with mono-ancestry parents (EAS × EUR), in which the entire genome is informative. The remaining 20 patients shared one ancestry across both haplotypes while carrying at least one distinct ancestry on a single haplotype. In these patients, the part of the genome with the shared ancestry cannot be attributed to either parent, while the haplotype-specific ancestry unambiguously identifies one parental chromosome. Only these parent-discriminant segments, spanning on average roughly a third (30.25%) of the genome, were retained for downstream analyses. Discriminant ancestries were predominantly EUR, EAS and SAS, reflecting their higher representation in the cohort, while AFR and EUR were the most frequently shared ancestries between haplotypes (**Figure 3A**).

**Figure 3.**
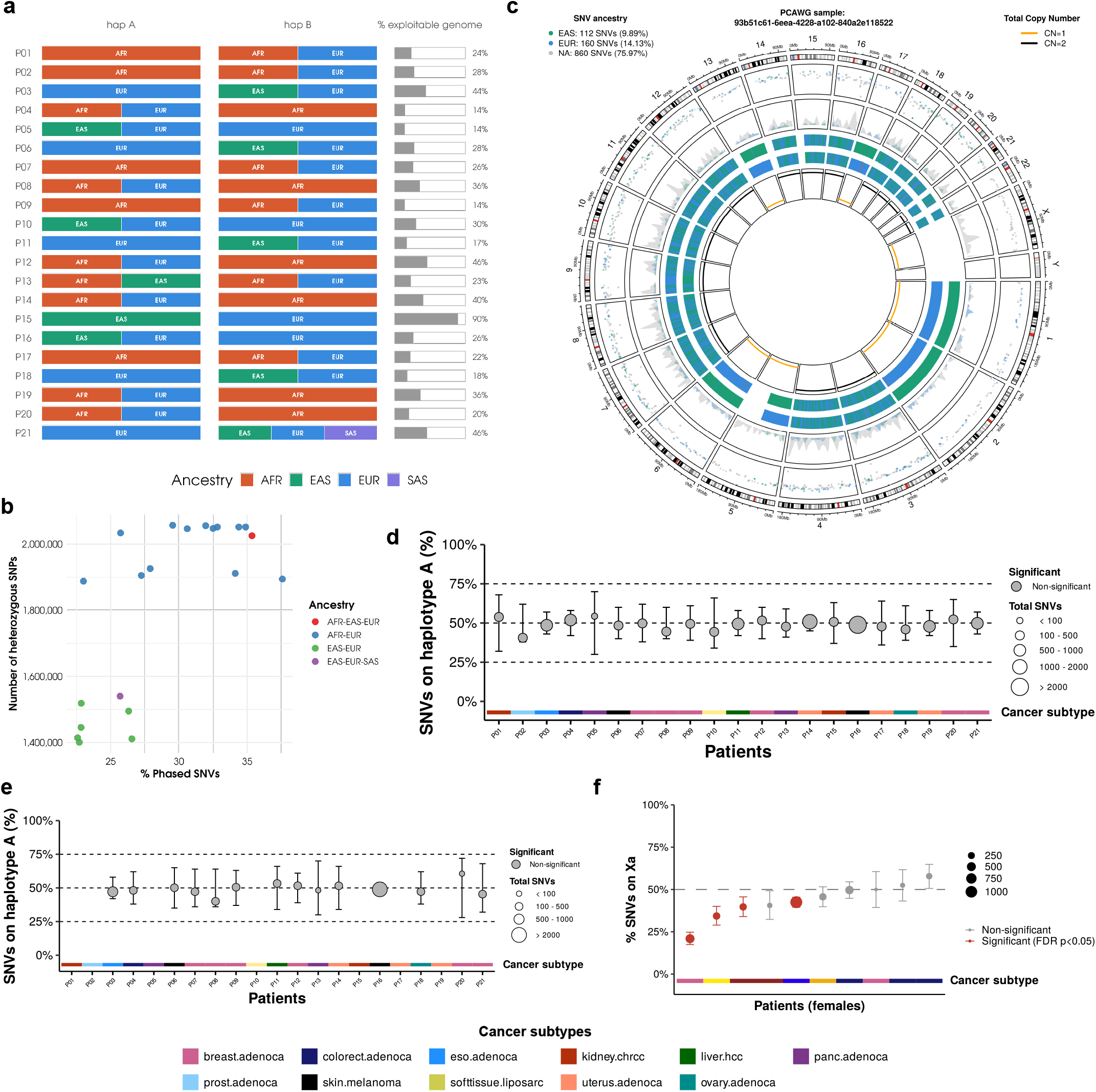
Very stable mutation accumulation across parental haplotypes in F1 admixed PCAWG genomes. **(A)** Local ancestry composition of hap A and hap B across the 21 informative patients (AFR, EAS, EUR, SAS), with percentage of parent-discriminant genome shown on the right. **(B)** Fraction of SNVs successfully phased relative to heterozygous SNPs. **(C)** Local ancestry profile of a representative informative patient inferred with Gnomix. Tracks (inner to outer): total copy number, haplotype-level local ancestry, ancestry-specific SNV density, and individual SNVs by ancestry. **(D-E)** Fraction of SNVs assigned to one haplotype accross informative patients with >50 SNVs. Each point represents a patient, sized by total SNVs; error bars show the maximum haplotype imbalance still consistent with each patient’s observed data (a = 0.05), accounting for phasing and local ancestry inference error. Dashed line at 50% indicates the expected equal distribution. Red points denote significant deviation from 50% after FDR correction (p < 0.05). **(D)** Allelic balanced regions; **(E)** Allelic imbalanced regions without loss of heterozygosity. **(F)** Fraction of SNVs on the active X chromosome (Xa) across female patients. Points are sized by total X-linked SNVs, with 95% binomial confidence intervals. Dashed line at 50% marks the expected distribution; red points indicate significant deviation after FDR correction (p < 0.05).

### Stable global mutation burden across parental genomes of different ancestries having coevolved within the same nuclei

We applied PhaSoMix to quantify allele-specific mutation rates in these 21 admixed PCAWG patients (**Supplementary Table 1**). A key factor limiting statistical power is the number of mutations that can be assigned to each parental haplotype (**Extended Data Figure S6**). In our 21 patients, the number of SNVs ranged from 1,086 to 91,690 (median 5,882). Somatic mutations were empirically phased using heterozygous SNPs present on the same sequencing read pairs, limiting analysis to mutations overlapping a heterozygous SNP. Among 23,166,019 candidate SNP positions, approximately 6.4% (∼1.48 million) were heterozygous, with a median inter-heterozygous SNP distance of 564 bp. Given paired-end 100 bp reads covering ∼400 bp at ∼55X coverage, a mean of 32.77% of SNVs per sample could be non-ambiguously phased. AFR-EUR admixed individuals exhibited higher phasing rates, reflecting the increased density of heterozygous SNPs driven by the differentiation between African and European haplotypes (**Figure 3B, Extended Data Figure S2 A-B**).

Each phased SNV was then assigned to a parental haplotype by querying the Gnomix local ancestry prediction at its genomic position (**Supplementary Figures 1-20**). Where the two haplotypes carried different ancestries, the mutation could be unambiguously attributed to one parent. To illustrate this, we examined the F1-admixed EUR-EAS female case with chromophobe renal cell carcinoma (**Figure 3C**). Local ancestry reconstruction confirmed an approximately 50% EUR and 50% EAS composition genome-wide (**Figure 3C**). In allelic-imbalance regions, where switch errors had been corrected using copy-number information, each haplotype was entirely mono-ancestral, with one haplotype exclusively EUR and the other exclusively EAS (**Figure 3C**). In balanced regions, alternating ancestry segments along a single phase reflected residual phasing switch errors, introducing noise in ancestry assignment. Of 2,029 autosomal SNVs detected in this patient, 567 (28%) could be phased, of which 191 were assigned to the paternal genome and 292 to the maternal genome. After correcting for copy number and removing loss of heterozygosity (LOH) regions, the paternal and maternal haplotypes harboured 104 and 101 somatic mutations, respectively. We quantified parent-of-origin-specific mutation burden across three genomic partitions: all regions combined (**Figures 3D, Extended Data Figure S7**), balanced regions (**Extended Data Figure S8**), and allelic-imbalance regions (**Figures 3E, Extended Data Figure S9**), where corrected phasing provides the most reliable ancestry assignment.

Across all 21 informative patients, mutation burdens in balanced regions were remarkably consistent between the two haplotypes, with all patients falling within the expected range under the null hypothesis of symmetric mutation accumulation (**Figure 3D**). To distinguish a true lack of asymmetry from limited statistical power, we asked, for each patient, how far the haplotype fraction could plausibly depart from parity given the observed data, after correcting for that patient’s own local ancestry and phasing accuracy (**Supplementary Table 3**). This upper bound was 4% in the best-powered patient (42 of 1,298 phased, assignable SNVs), and 5.7% when combining evidence across the entire cohort (12,839 assignable SNVs; binomial tests, Fisher’s method, p = 0.329). A complementary power analysis confirmed that, given the observed mutation burden and per-patient accuracy, the cohort had 90% power to detect a systematic asymmetry of 5% or greater, ruling out anything but small undetected effects. Finally, to test directly for a bias toward a specific parent rather than asymmetry of unknown direction, we examined the subset of 10 patients with resolved parental haplotype assignment: no directional bias toward either the paternal or maternal genome was observed (weighted effect: −0.03%, 95% CI: −7.25% to 2.22%; sign test, p = 0.34).

Together, these results indicate that a potential global asymmetric mutational pressure or repair between parental haplotypes would be very limited or rare, and that somatic mutation accrual is largely symmetric between the two parental genomes across cancers.

We further assessed functional genomic regions, including genes, intergenic regions, introns, and exons, and observed consistent SNV burdens between the two parental alleles across region types (**Extended Data Figure S7-9**). For some patients, the number of phased mutations within specific regions was too low to draw reliable conclusions. A small number of patients showed statistically significant differences in particular regions, but these were isolated observations and the large majority showed no imbalance, consistent with the genome-wide result.

Imprinted regions are characterized by local parent-of-origin-specific epigenetic asymmetry between maternal and paternal alleles, making them a relevant context in which to investigate allele-specific mutation rates. We investigated imprinted loci using the established catalogue of imprinted regions^20^. However, only 2 mutations were detected and phaseable in these regions across the 21 patients, precluding meaningful analysis of allele-specific mutation rates in imprinted loci. As another positive control, we next investigated the X chromosome in female patients, and assessed whether the inactive copy exhibits differential mutation accumulation. We quantified the number of mutations occurring on the active and inactive X chromosomes in 11 female patients from the PCAWG cohort. In 8 out of 11 patients, a higher number of mutations was observed on the inactive X chromosome, with 4 patients showing a statistically significant difference between the two copies (**Figure 3F**), consistent with prior observations^21^.

### Stable mutation burdens in the clonal and subclonal periods, and at (C>T)pG

Copy-number gains, frequent events in cancer genomes, are effectively molecular snapshots in time of the gained copy. This is because at sequencing, mutations present on the two copies (m2) accumulated over time between the fertilized egg and the time of the gain, while mutations on one of the copies (m1) accumulated over time after the gain and until resection^5^. Using this rationale, not only can mutations be timed before (m2 mutations) or after (m1 mutations) the gain, together m2 and m1 also allow to derive molecular timing estimates of the gains, which can be integrated across samples into full cancer timelines^4^. Akin to most cancer genomics analyses, copy-number gain-timing generally assumes that the two parental copies, as well as copies of the parental homologs, accumulate somatic mutations at similar rates. To evaluate the consequence of violations of this assumption, we measured how increasing differences in mutation burden between parental haplotypes affect gain-timing accuracy. As the mutation imbalance grows, the inferred gain time progressively drifts away from the value expected under equal mutation rates, showing a clear bias driven by unequal haplotype mutation loads (**Figure 4A**). Importantly, this bias becomes stronger for gains that occurred earlier in tumor evolution, with the estimation error increasing steadily as the true gain time rises (**Figure 4A**).

**Figure 4.**
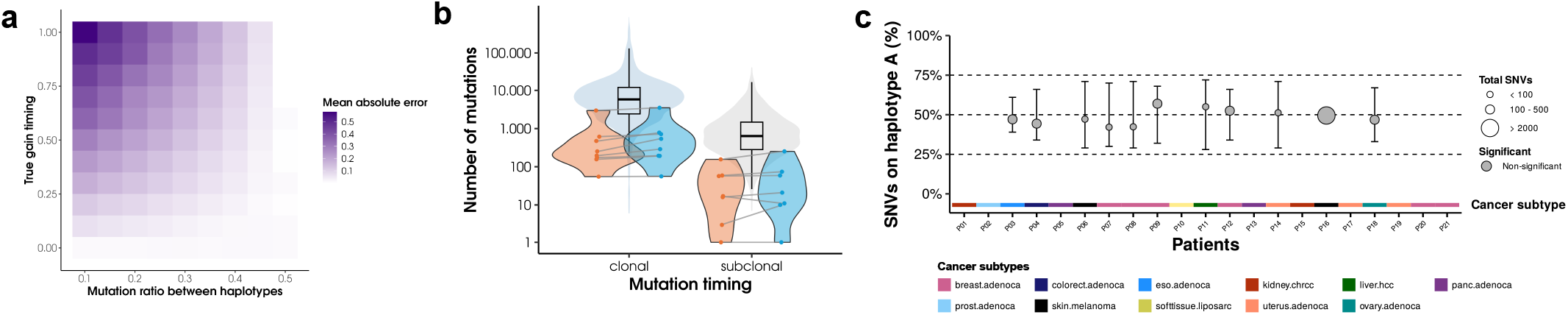
Stable mutation burdens in the clonal and subclonal periods, and at (C>T)pG. **(A)** Effect of parental mutation imbalance on gain timing estimation. The x-axis represents the proportion of mutations assigned to one haplotype (e.g., 0.3 indicates 30% of mutations on haplotype A and 70% on haplotype. The y-axis shows the true gain timing. Color intensity reflects the mean absolute error between the predicted and true gain timing across 100 simulations. **(B)** Background: Distribution of mutations across gain timing categories (clonal, and subclonal) for the entire PCAWG cohort. Forground: counts of clonal and subclonal mutations for each parental haplotype across admixed PCAWG patients. Each dot represents one haplotype in a patient. **(C)** Fraction of C>TpG SBS1 SNVs assigned to the maternal haplotype across F1 patients with >50 SNVs in alleleic imbalanced regions. Each point represents a patient, sized by total SNVs; error bars show the maximum haplotype imbalance still consistent with each patient’s observed data (α = 0.05), accounting for phasing and local ancestry inference error. Dashed line at 50% indicates the expected equal distribution. Red points denote significant deviation from 50% after FDR correction (p < 0.05).

Moreover, even if the overall mutation load remains comparable between the two parental copies, mutations could in principle have accumulated at different times on the maternal and paternal haplotypes, leading to temporal differences in mutation rates. To address this possibility, we quantified, for each admixed patient, the number of clonal and subclonal mutations assigned to each parental haplotype. Our results show that maternal and paternal copies accumulate mutations in similar proportions across timing categories, with no statistically significant differences, showing that there are also no detectable asymmetry in those time windows (**Figure 4B**).

The susceptibility of methylated CpG sites to spontaneously deaminate leading to (C>T)pG transitions (SBS1 clock-like signature) ^22^ could in theory lead to differences in mutation rates between parental genomes with different CpG methylation rates in some individuals. Yet, we observed no allelic bias in (C>T)pG transitions genome-wide, indicating that clock-like mutation accumulation remains symmetric across parental alleles (**Figure 4C**).

## Discussion

In cancer genomics analyses, such as mutation calling, driver identification, or gain timing, the two homolog parental copies are implicitly assumed to accrue mutations at the same rates. Given our current understanding of somatic mutation accrual and repair, although this assumption seems reasonable especially at the genome-scale, it has never been tested in cancer datasets. Also, it is unknown how effects of violation of this assumption could impact the results of such analyses, i.e. mutation calls, the identification of cancer driver genes, or molecular timing estimates.

Multiple factors, including sequence, epigenetic marks, levels of expression, and chromatin structure, are known to influence local mutation rates in an allele-specific manner^12,23^. However, it is still unclear how such factors could translate to more global or genome-wide differences in mutation rates between the two parental copies. While the factors and mechanisms for such a global difference would be speculative at this point, it can be reasoned that such factors might exist. For example, parental environmental conditions such as obesity or starvation have been shown to induce genome-wide changes in CpG methylation patterns ^14,24–26^, that could be maintained after epigenetic reprogramming and even transmitted across generations outside imprinted regions. Although the inheritance is not necessarily always mediated by surviving epigenetic marks, in theory, any parent-of-origin differential in CpG methylation leakage after the epigenetic resets is expected to lead to differences in (C>T)pG mutation rates on the affected parental genome in the offspring.

To answer this question, we developed PhaSoMix, which harnesses the high genetic distance between parental genomes of mixed-ancestry patients to phase their genomes to the parent/ancestry of origin. PhaSoMix identified 21 recently-admixed patients out of the 2,645 patients in the PCAWG dataset. After quantifying the mutation rates on each parental copy, remarkably stable mutation burdens are observed, even at (C>T)pG in all individuals and across cancer types. Thus, this novel approach validates that parental genomes mutate at similar global rates, an important assumption in cancer genomics analyses, including gain timing and molecular archaeology^5^. Thanks to our framework to propagate errors, we can derive an upper bound for the differences in mutation burden. Leveraging this upper bound to model its effect on timing estimates, we demonstrate that they would be minimal in most time windows.

PhaSoMix is conceptually related to surrogate-parent phasing^27^, but uses ancestries rather than relatives as surrogate parents: it leverages haplotype-frequency differentiation between super-populations rather than identity-by-descent between family haplotypes. Access to the genotypes of relatives would likely become much more potent for phasing, and more feasible as large genotype databases grow in the future. Meanwhile, *PhaSoMix* can be applied to cancer samples without such large population-wide databases, which are restricted in their access, and in individuals for which the two parental copies already are genetically distant.

There are limitations to our study. First, PhaSoMix can only phase admixed individuals, a small fraction of the population, and thus requires a large cancer genome database to identify enough such cases. Second, while different ancestries maximize genetic distance between the parental copies to allow genomewide phasing, this genetic distance also increases the local differences in mutation rates^28^, which could hide more subtle genomewide signals. While ancestry-based phasing is limited, PhaSoMix modules for SNP-in-read phasing and copy-number-aware quantification on each parental haplotype have broad applicability.

In the future, growing cancer genome databases and adoption of long-read sequencing, coupled with large genotype reference sets would allow us to phase more cancer (epi)genomes to their parent of origin^16,27,29^, potentially revealing cases of significant global differences between (epi)mutation rates of the two parental genomes in specific clinical contexts.

## Material and methods

### 1000 Genomes and Pan-Cancer Analysis of Whole Genomes data

We downloaded both the Phase 3 low-coverage 1000 Genomes Project (1kGP) VCFs (GRCh37, 2,504 unrelated individuals)^30^ and the recent high-coverage 1kGP dataset (GRCh38, 3,202 samples including 602 trios)^31^. To create a high-coverage GRCh37 resource, we matched SNPs by their IDs and retained only those present in both datasets, effectively merging the two to obtain a harmonized set of high-confidence genotypes in GRCh37.

BAM files for the Pan-Cancer Analysis of Whole Genomes (PCAWG)^32^ were sourced from the Genomic Data Commons (GDC) and the European Genome-Phenome Archive (EGA) and processed with alleleCounter v4.2 and the first step of the Battenberg pipeline v2.2.1^33^ to generate VCF files. Within these steps, phasing is conducted with Beagle 5.4^34^ and the 1kGP GRCh37 reference map. For both datasets, we filtered out variants that were not single nucleotide polymorphisms (SNPs) and absent in either dataset.

### Global ancestry inference

Admixture fractions were estimated using Admixture^35^, with the 1000 Genomes Project (1kGP) dataset serving as the reference panel. For the training, only unrelated individuals from 1kGP were retained, and models were trained at two different levels, Super-Populations and Populations (limited to their respective Super-Populations). SNPs with minor allele frequency (MAF) below 0.1 were removed, and each SNP was assigned a score reflecting its discriminative power (Formula 1). The analysis focused on the 100,000 most discriminative SNPs, ensuring balanced representation across populations. These SNPs were used as input for unsupervised ADMIXTURE analyses.

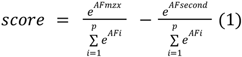

Patients were considered of a single ancestry when their dominant ancestry proportion exceeded 0.95. These individuals were incorporated into the Gnomix training set alongside 1kGP samples, increasing haplotype diversity and the number of training individuals available for local ancestry inference.

### SNV phasing

Heterozygous SNVs were empirically phased to the nearest heterozygous SNP using raw sequencing data. For each individual sample, Samtools mpileup (v1.21) was run on the corresponding BAM files, and the mpileup output was parsed to extract read Q-names at both the heterozygous SNP and heterozygous SNV positions. Matching Q-names at genotype-position allowed to quantify reads simultaneously carrying a SNP reference or alternate allele and a SNV alternate allele. Any SNV showing ambiguous phasing, specifically, cases where reads linked a SNV to both the reference and alternate alleles of a heterozygous SNP, was excluded and represented only 0.95% of phasable SNVs. All non-ambiguous SNVs were retained, including those phased to a single informative SNP, ensuring maximal inclusion of variants while maintaining phasing accuracy.

### Local ancestry inference

Local ancestry inference was performed using Gnomix (window size 0.1 cM) trained on phased haplotypes from 1kGP and single ancestry PCAWG individuals. Seven model sets were trained: one at the super-population level (AFR, AMR, EUR, SAS, EAS) and one per super-population resolving its constituent sub-populations. Sub-population models showed insufficient accuracy and were discarded. All analyses were therefore performed using the super-population model. Each phased SNV was assigned the ancestry of its carrying haplotype at that genomic position.

### Identification of informative patients

Local ancestry was inferred per haplotype window with Gnomix. For each ancestry pair, we computed the heterozygous fraction within somatic copy-number imbalance zones (IZ; major ≠ minor copy number, length > 10 Mb). Patients with a dominant heterozygous pair covering ≥90% of IZ windows and <1% homozygosity were classified as strict F1. Patients with 20–90% coverage and the same homozygosity threshold were classified as partially informative. In these patients, an ancestry was retained as discriminant if heterozygous in >5% of the genome, homozygous in <10% of occurrences, and homozygous on both haplotypes in at most two IZ regions.

Parental haplotypes were resolved by graph bipartition. Ancestry pairs co-occurring in heterozygous IZ windows were linked in a graph, weighted by shared base pairs. Ancestries homozygous on both haplotypes in >5% of IZ base pairs were assigned to both haplotypes. The remaining graph was 2-coloured by breadth-first search. Bipartite components yielded direct haplotype assignments; non-bipartite components were resolved at the biconnected-component level, with unresolved ancestries flagged as ambiguous.

### SNV quantification correction

In regions with equal numbers of each parental copy (balanced regions), SNV are quantified by counting the number of SNVs attributed to each ancestral haplotype. In contrast, imbalanced regions, where one parental allele has more copies than the other, require an adjusted approach to account for the accrual of somatic mutations due to the increased number of copies. In these imbalanced regions, the multiplicity of each SNV is divided by the copy number of the corresponding parental copy carrying the variant (Formula 2). This correction ensures that the influence of varying copy numbers is appropriately factored into the analysis.

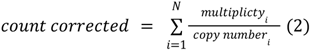

### Paternal and maternal ancestry inference

The ancestry of each SNV is assigned, but the parental origin of these variants must still be resolved. Maternal ancestry is determined using mitochondrial DNA (mtDNA). In males, maternal and paternal origins can be further validated through ancestry inference of non-paralogous regions of the X and Y chromosomes, respectively. For each mixed-ancestry PCAWG patient, mtDNA and informative chrX/chrY loci were genotyped from the normal BAM using AlleleCounter v4.3.0 (https://github.com/cancerit/alleleCount) to obtain allele counts and infer genotypes. To assign parental origin, XGBoost models were trained separately for chromosome X, chromosome Y and mtDNA using SNPs from 1kGP haplotypes, with the four super-populations (AFR, EUR, SAS, EAS) as class labels and a softmax objective. For regions containing more than 100,000 SNPs, only the 100,000 most variable SNPs across training haplotypes were retained. Each informative patient was classified using the corresponding chromosome-specific model. Parental ancestry was manually assigned to each haplotype only when the predicted probabilities provided an unambiguous distinction between the possible ancestry combinations of the two haplotypes.

### Allele-specific assignment of somatic mutations to the active and inactive X chromosomes

We analyzed 11 female tumors from the Pan-Cancer Analysis of Whole Genomes selected for high SNV burden and extensive allelic imbalance on chromosome X. Imbalance zones (major_cn ≠ minor_cn; minor_cn > 0) were defined from consensus copy-number segments. Within each region, heterozygous SNPs displaying allelic expression bias in RNA-seq data (coverage ≥10, allelic ratio ≥0.65), excluding pseudoautosomal regions, were used to infer the transcriptionally active X chromosome (Xa): for each SNP, the overexpressed RNA allele was matched to DNA allele-specific counts to determine whether Xa corresponded to the major or minor copy, and a weighted vote across SNPs (minimum 70% concordance) assigned Xa at the segment level. Somatic SNVs were phased to nearby heterozygous SNPs, allowing each mutation to be assigned to the gained or non-gained haplotype. By combining this information with the segment-level identification of the active X chromosome, we classified each mutation as occurring on Xa or Xi. Mutation counts were adjusted for local copy number and mutation multiplicity, and deviation from a 1:1 Xa/Xi ratio was tested using false discovery rates from two-sided binomial tests.

### Switch error correction

Phasing errors in allelic-imbalance segments were corrected in two steps to generate a local “ground truth” for phasing accuracy (**Extended Data Figure S3)**. These regions are particularly informative because copy-number differences between parental haplotypes create detectable deviations in B-allele frequencies (BAF), providing strong signals to identify and correct phasing errors. First, contiguous haplotype blocks showing consistent BAF deviations were identified using piecewise-constant fitting and flipped if necessary. Second, individual SNPs were further refined using a likelihood-ratio test comparing observed BAF to expected values under each haplotype configuration. By correcting switch errors in these allelic-imbalance regions, we obtain a reliable reference that allows quantitative assessment of phasing accuracy. The switch error (SE) rate was calculated as the proportion of SNPs whose phase had to be inverted relative to the original Beagle output.

### Statistical framework for haplotype imbalance

Local ancestry inference and phasing errors attenuate the true haplotype ratio toward 0.5 in a predictable way; all analyses correct for this using each patient’s local ancestry and phasing accuracy. For each patient, we determined the largest haplotype imbalance, in the observed direction, compatible with the observed counts (exact binomial test, α = 0.05). Per-patient p-values were combined across the cohort via Fisher’s method to obtain both a cohort-level significance test and an equivalent compatibility bound. Power was estimated as the probability that a given imbalance, present in every patient, would yield at least one FDR-significant patient (5,000 simulated replicates, using each patient’s mutation burden and accuracy). In patients with parental haplotype assignment, per-patient paternal fractions were corrected for pipeline accuracy and combined via a burden-weighted bootstrap (5,000 resamples) and an exact sign test.

### Algorithm testing and validation

#### Local ancestry inference

Accuracy was evaluated using simulated admixed individuals generated from a held-out 20% test set of 1kGP haplotypes. Mixed-ancestry individuals were constructed by combining haplotypes from two unrelated non-admixed individuals, with switch errors introduced at varying frequencies to simulate phasing noise **(Extended Data Figure S10)**. For each simulated individual, the true ancestry of every 0.1 cM segment was known by construction. Gnomix models trained on the remaining 80% were used to infer local ancestry. For each segment, each haplotype was scored as correct (1) or incorrect (0), and the two scores averaged to give a per-segment accuracy of 0, 0.5, or 1. Accuracy was reported separately for each ancestry combination and SER level..

#### SNV phasing

SNV phasing was validated using two complementary approaches. First, some SNVs can be phased to multiple nearby SNPs carried by different reads, thereby we can assess the consistency across SNPs from the same haplotypes. We count a correct phase when all the SNPs from the same haplotype phase consistently with the SNVs. Another method involves looking at the number of SNVs in a segment with allelic imbalance. SNVs for which the inferred multiplicity is higher than the number of minor alleles should be phased to the major allele. This is particularly informative in regions with clonal loss-of-heterozygosity (LOH), where SNVs should never be phased to the lost allele.

#### SNV quantification correction

The validity of the copy-number correction for SNV quantification was assessed through three complementary approaches. First, using simulated data with known mutation rates, copy numbers, and copy-gain timings, we compared the estimated mutation rate (with and without correction) to the true simulated rate. Second, in imbalanced segments of mixed-ancestry PCAWG patients, we tested whether the mutation rate measured from mutations phased to gained copies for one allele with CN > 1 matched the reference rate measured from mutations phased to the other allele with CN = 1 across allelic ratios (e.g., 1:2, 1:3, 1:4, 1:5), before and after correction for the number of copies. Third, across all PCAWG samples, we compared mutation rates measured from mutations in gained segments with loss of heterozygosity (LOH) with mutation rates measured from mutations in segments with a single copy of the major allele, before and after correction for the number of copies.

#### Paternal and maternal ancestry inference

Parental ancestry inference of sex chromosome and mitochondrial variants was assessed using VCFs, generated from 1kGP BAM files, split into 80/20% train/test sets for every pairwise combination of ancestral populations. Analyses of chrX and chrY were restricted to non-paralogous regions. Population and subpopulation reference labels derived from 1kGP metadata were used to run XGBoost separately for mtDNA, chrX, and chrY, producing per-individual ancestry proportions for each ancestry. Predicted ancestries were compared to true labels to assess the accuracy.

#### SNP Phasing

Informative admixed PCAWG genomes were phased using Beagle 5.4^34^. Because accurate SNP phasing is critical for reliable ancestry inference of somatic mutations in admixed genomes, we tested different reference panels, which can influence phasing accuracy. We evaluated three distinct panels: (i) the low-coverage 1000 Genomes Project panel in GRCh37, (ii) a high-coverage 1000 Genomes Project panel originally in GRCh38 and lifted over to GRCh37 to match PCAWG VCFs, and (iii) the high-coverage panel further extended with non-admixed PCAWG individuals whose genomes had been manually corrected in allelic-imbalance segments.

## Data availability

The data used in this study are publicly available. Genomic reference data were obtained from the 1000 Genomes Project^30^ (https://ftp.1000genomes.ebi.ac.uk/). The data used in this study are publicly available through the ICGC/TCGA Pan-Cancer Analysis of Whole Genomes (PCAWG) Consortium. Whole-genome sequencing BAM files for ICGC samples are available via the European Genome-phenome Archive (EGA; EGAS00001001692) under controlled access. BAM files for TCGA-derived samples are available via Bionimbus/ICGC (https://icgc.bionimbus.org/files) under controlled access. Somatic variant calls for ICGC samples can be accessed via the ICGC ARGO platform (https://platform.icgc-argo.org/), and variant calls for TCGA samples are available via Bionimbus/ICGC (https://icgc.bionimbus.org/files).

## Code availability

The code to reproduce the analyses in this study as well as run PhaSoMix is available on Github: https://github.com/IRIBHM-computational-groups/PhaSoMix, and the version used in this study has been deposited on Zenodo: https://zenodo.org/records/21495717.

## Acknowledgements

This work was supported by the Fonds de la Recherche Scientifique - FNRS under Grant(s) n° 40017793. PVL is a CPRIT Scholar in Cancer Research and acknowledges CPRIT grant support (RR210006). This work used data generated by the TCGA Research Network (https://www.cancer.gov/tcga), the ICGC/TCGA Pan-Cancer Analysis of Whole Genomes Consortium and the 1000 Genomes Project Consortium. We gratefully acknowledge the contributions of the patients, volunteers, and research groups who provided these data. We thank members of the IRIBHM computational labs, Jonas Demeulemeester, Nil Fernandez Lojo, Guillaume Smits and Isabelle Migeotte for their valuable input.

## Contributions

ML conceived the experiments, developed PhaSoMix, ran and co-supervised bioinformatics analyses, and wrote the manuscript. AC ran bioinformatics analyses. MP developed the SNP-in-read SNV phasing code. PVL & VD provided guidance and conceptual input. MT conceived the experiments, supervised the work, and wrote the manuscript. All authors approved the manuscript.

---

**Extended Data Figure S1.**
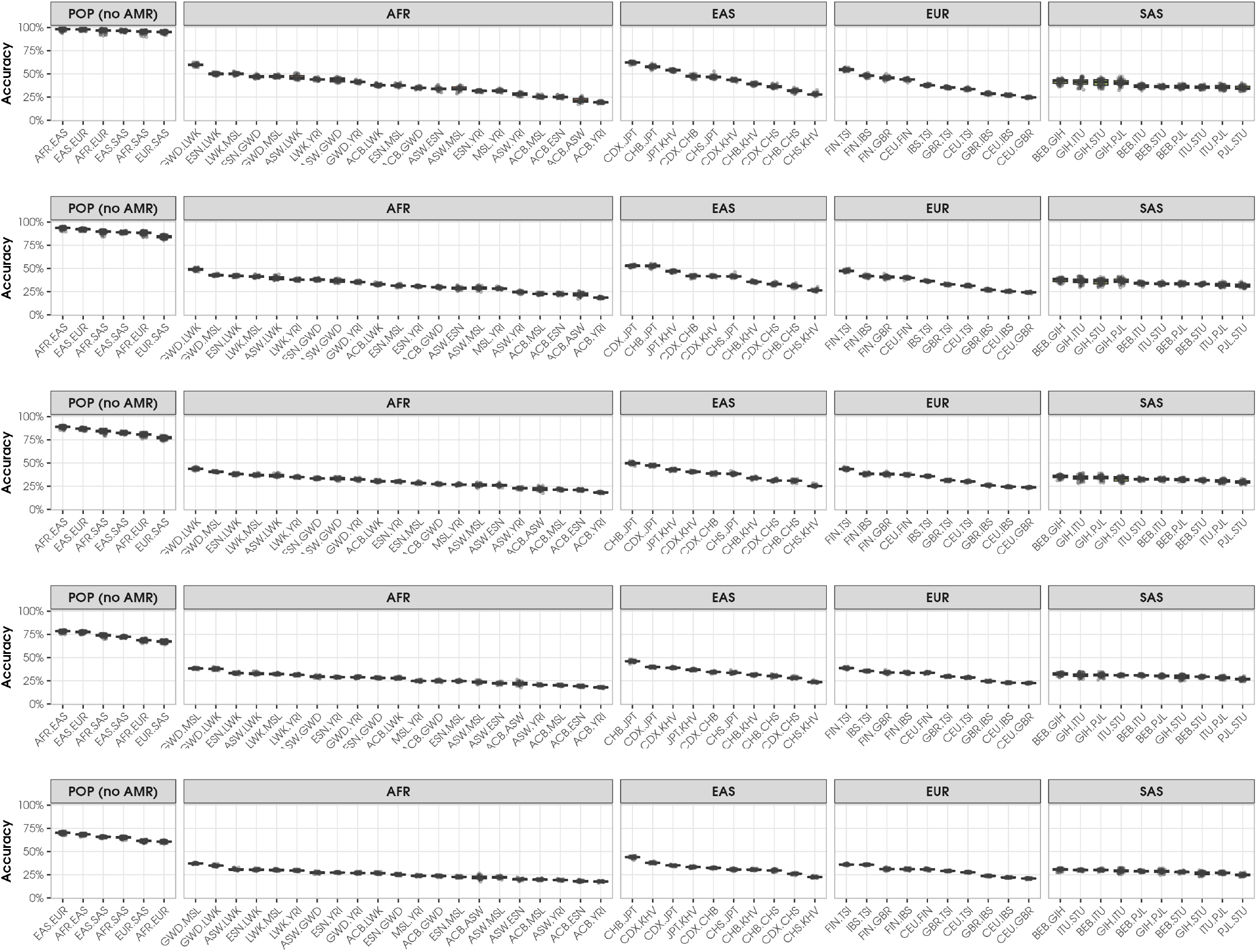
Chromosome-pair accuracy of local ancestry inference (Gnomix) across increasing switch-error rates. Panels correspond to switch-error rates (SER) of 0%, 0.1%, 0.2%, 0.5%, and 1%. Within each SER panel, sub-panels correspond to a super-population (AFR, EAS, EUR, SAS) or the full population set excluding AMR (POP no AMR). Boxplots show the distribution of weighted mean accuracy across chromosomes 1-22 for each ancestry pair combination, ordered by decreasing median accuracy. Individual points represent replicate simulations.

**Extended Data Figure S2.**
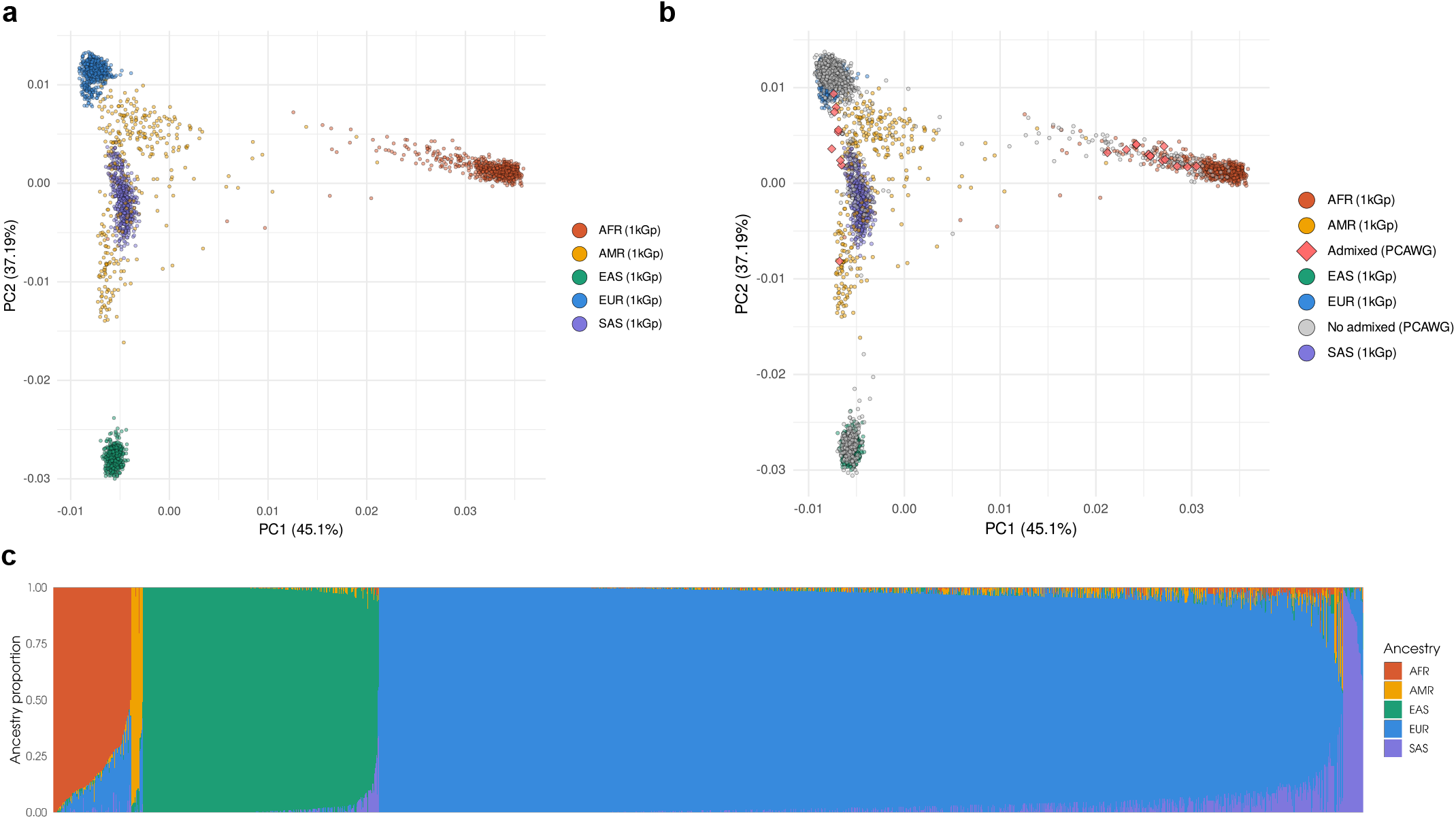
Genetic ancestry structure of PCAWG cancer genomes inferred from 100,000 most discriminant SNPs. **(A)** Principal component analysis on 100.000 most discriminants SNPs of 1kGp super-populations. **(B)** Principal component analysis on 100.000 most discriminants SNPs of 1kGp super-populations and admixed PCAWG patients. 1kGp points are colored by self-reported ancestry. Axes indicate variance explained by the first two PCs. **(C)** Global ancestry proportions of PCAWG patients estimated by supervised ADMIXTURE. Each vertical bar represents one patient; colors indicate the contribution of each super-population ancestry (AFR, AMR, EAS, EUR, SAS).

**Extended Data Figure S3.**
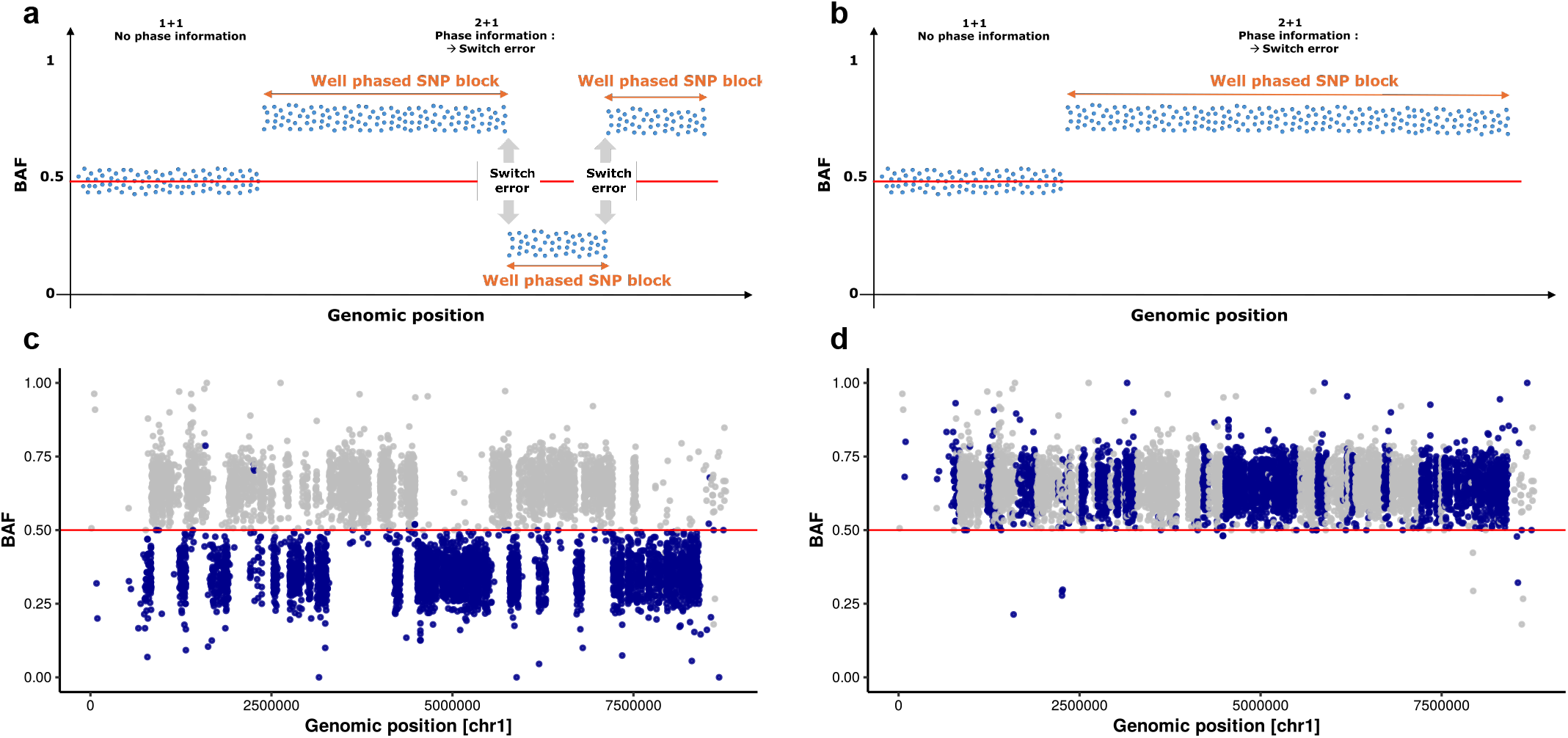
Phasing correction in allelic imbalanced segments. **(A-B)** Schematic representation of the phasing correction strategy in allelically imbalanced genomic segments. Each dot represents a heterozygous SNP, and the y-axis corresponds to the B-allele frequency (BAF) of one haplotype. Two types of segments are shown: (i) a balanced segment with equal copy number on both parental haplotypes, characterized by BAF values centered around 0.5; and (ii) an allelically imbalanced segment with a copy-number state of 2:1, for which the expected BAF values are ∼0.33 and ∼0.66 across the entire segment. **(A)** phasing switch errors are visible as abrupt transitions between the two expected BAF states (from ∼0.66 to ∼0.33 or vice versa). **(B)** Switch errors are corrected by enforcing phase consistency across the segment, i.e. by flipping the BAF value (i.e. 0.33 0.66) and haplotypes (i.e. 0|1 1|0) when necessary. (C-D) Example of BAF profiles of an alleleic imbalanced segment in chromosome 1 from a randomly selected admixed PCAWG patient. Stich of color represent switch errors. **(C)** Raw BAF signal prior to correction. **(D)** BAF after phasing correction.

**Extended Data Figure S4.**
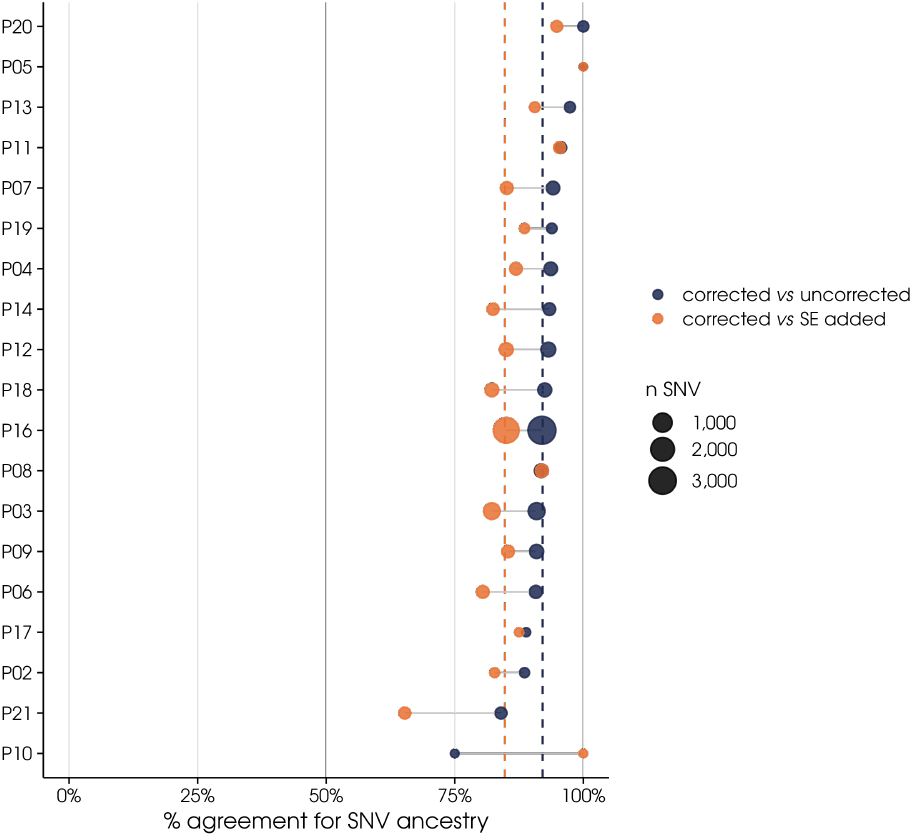
Per-patient SNV-weighted concordance of ancestry-based SNV assignments in allelic-imbalance regions. Corrected vs uncorrected phasing (real data) and corrected vs corrected+simulated switch errors are compared. Point size scales with SNV count; dashed lines show the weighted mean per comparison.

**Extended Data Figure S5.**
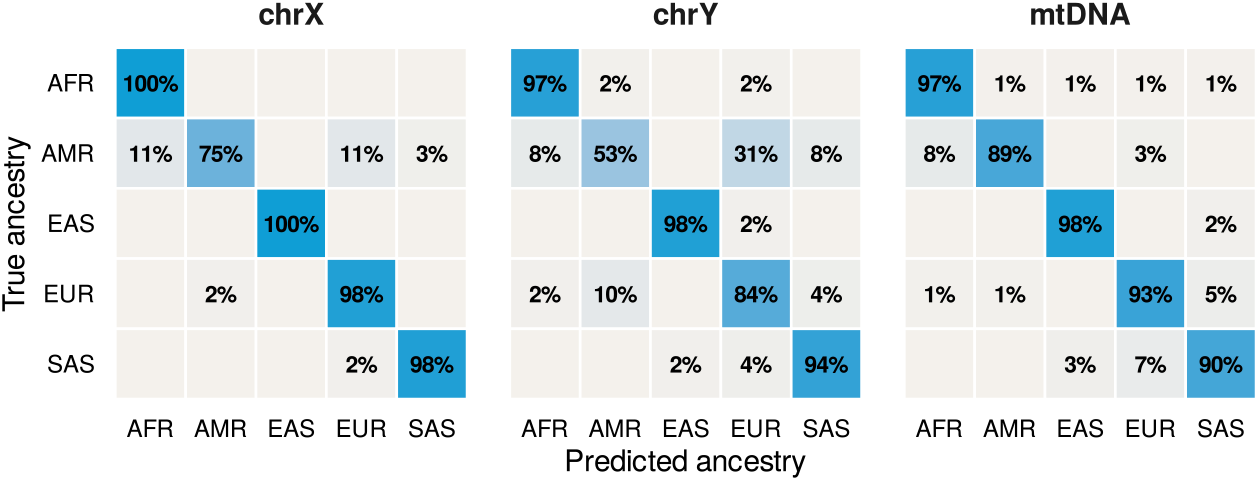
Ancestry inference accuracy on sex chromosomes and mitochondrial DNA. Confusion matrices for XGBoost-based parental origin classification across chrX, chrY, and mtDNA. Each cell shows the proportion of samples with a given true ancestry (rows) assigned to each predicted ancestry (columns), normalized by true class. Super-populations follow the 1000 Genomes Project classification (AFR, AMR, EAS, EUR, SAS).

**Extended Data Figure S6.**
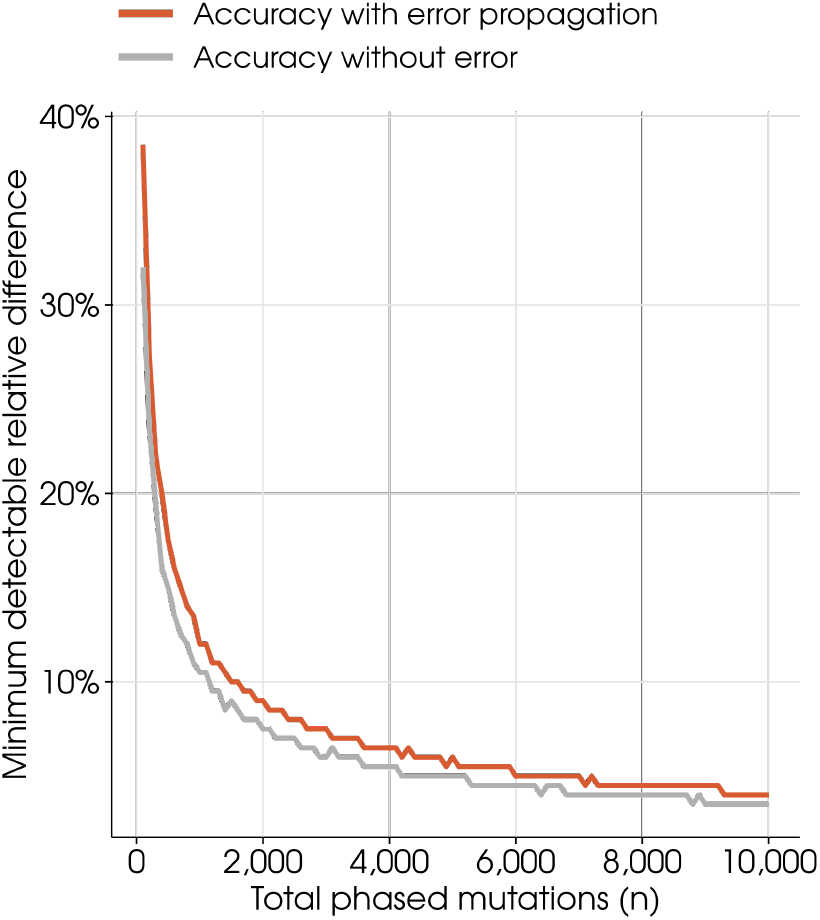
Statistical power to detect significant differences between parental copies. The minimum detectable relative difference in SNV counts between the two parental copies (90% power, α = 0.05) is shown across increasing total SNV counts. The orange curve incorporates pipeline error propagation (phasing and local ancestry inference accuracy); the grey curve represents the idealized case of perfect assignment accuracy.

**Extended Data Figure S7.**
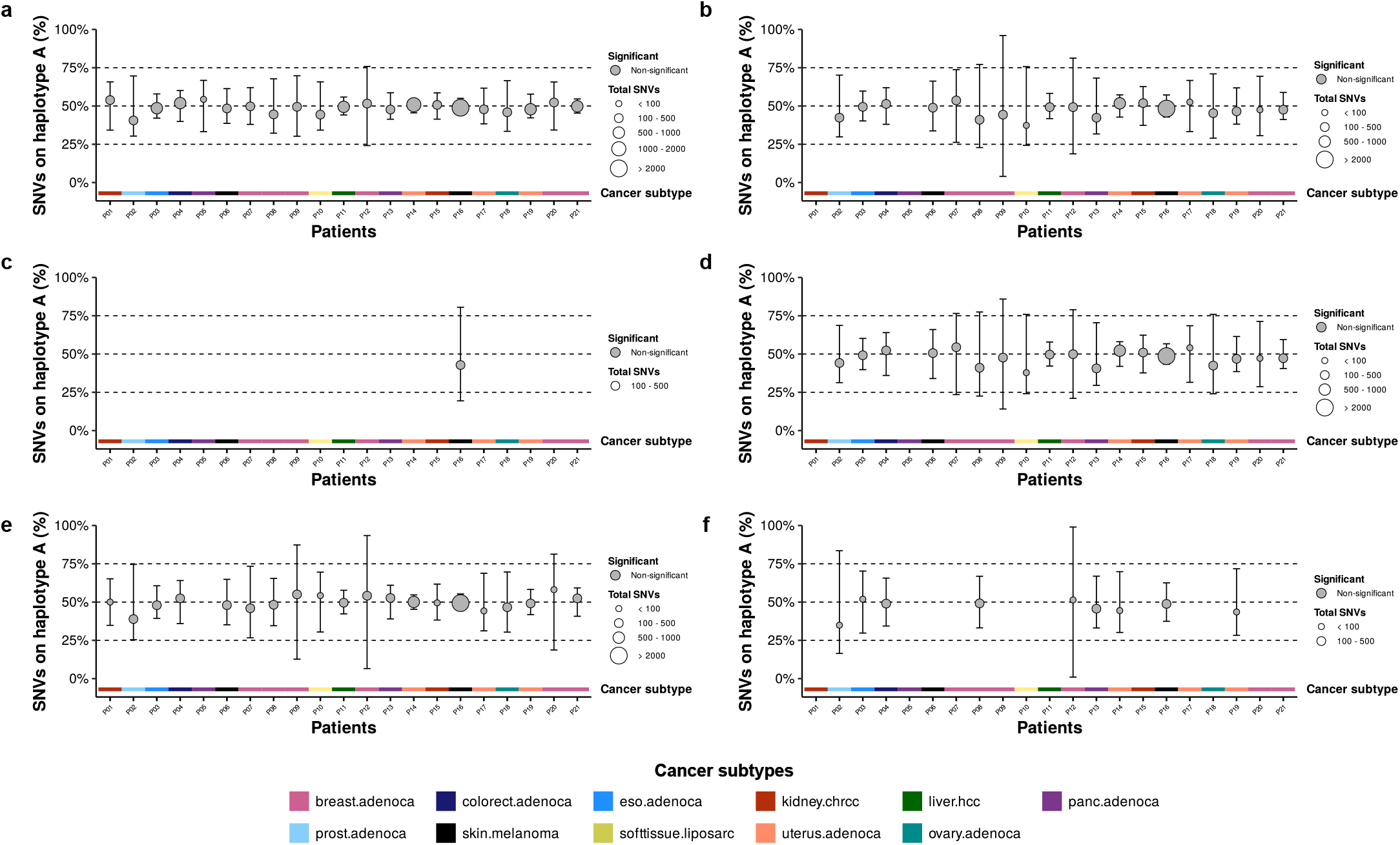
Very stable mutation accumulation across parental haplotypes in admixed PCAWG genomes. Fraction of SNVs assigned to one haplotype across admixed patients with ≥50 SNVs. Each point represents a patient, sized by total SNVs; error bars show the maximum haplotype imbalance still consistent with each patient’s observed data (α = 0.05), accounting for phasing and local ancestry inference error. Dashed line at 50% indicates the expected equal distribution. Red points denote significant deviation from 50% after FDR correction (p < 0.05). **(A)** Genome-wide (autosomes) **(B)** Genes **(C)** Exons **(D)** Introns **(E)** Intergenic regions **(F)** C>TpG (SBS1) SNVs.

**Extended Data Figure S8.**
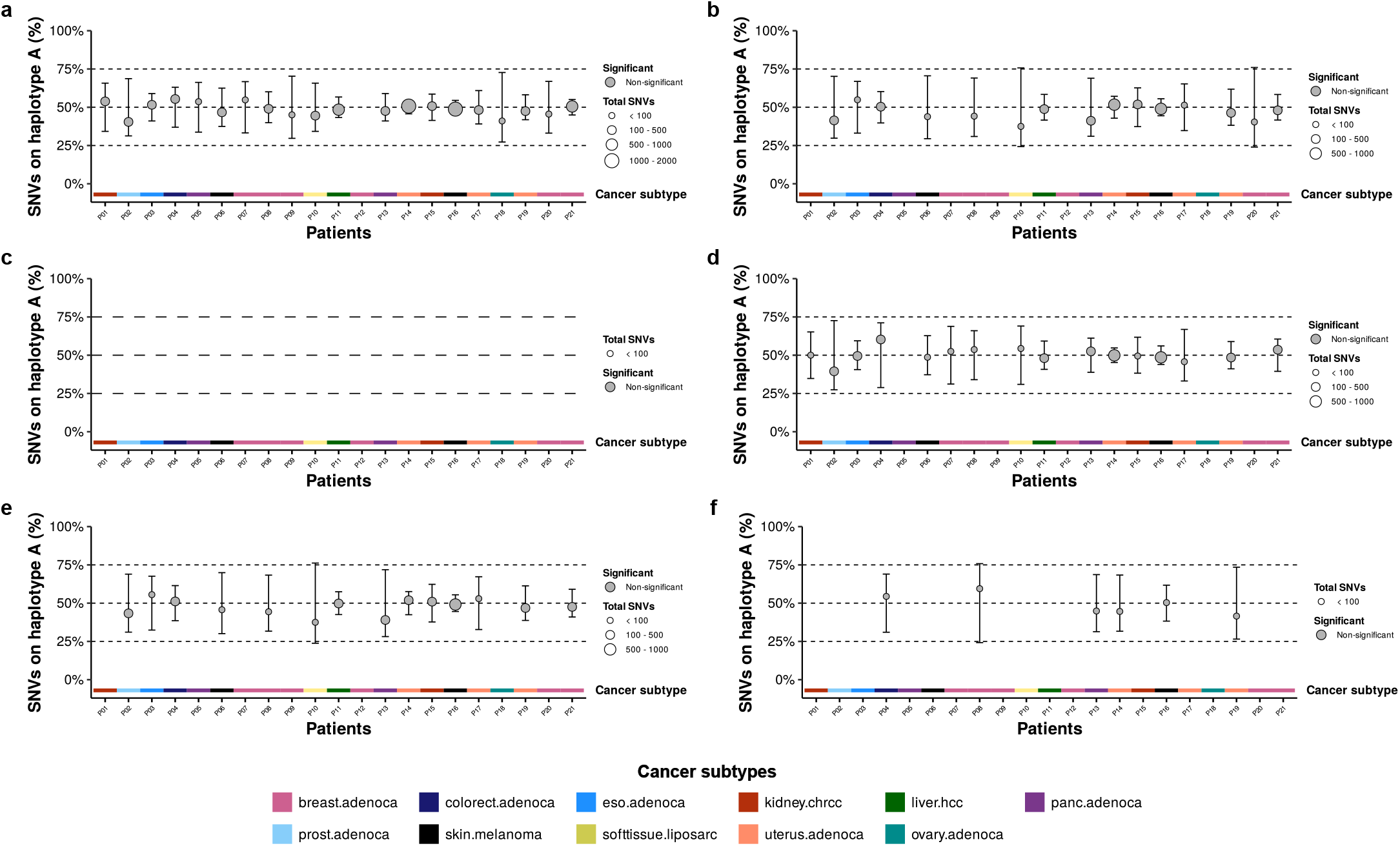
Very stable mutation accumulation across parental haplotypes in admixed PCAWG genomes in allelic balanced regions. Fraction of SNVs assigned to one haplotype across admixed patients with ≥50 SNVs. Each point represents a patient, sized by total SNVs;error bars show the maximum haplotype imbalance still consistent with each patient’s observed data (α = 0.05), accounting for phasing and local ancestry inference error. Dashed line at 50% indicates the expected equal distribution. Red points denote significant deviation from 50% after FDR correction (p < 0.05). **(A)** Genome-wide (autosomes) **(B)** Genes **(C)** Exons **(D)** Introns **(E)** Intergenic regions **(F)** C>TpG (SBS1) SNVs.

**Extended Data Figure S9.**
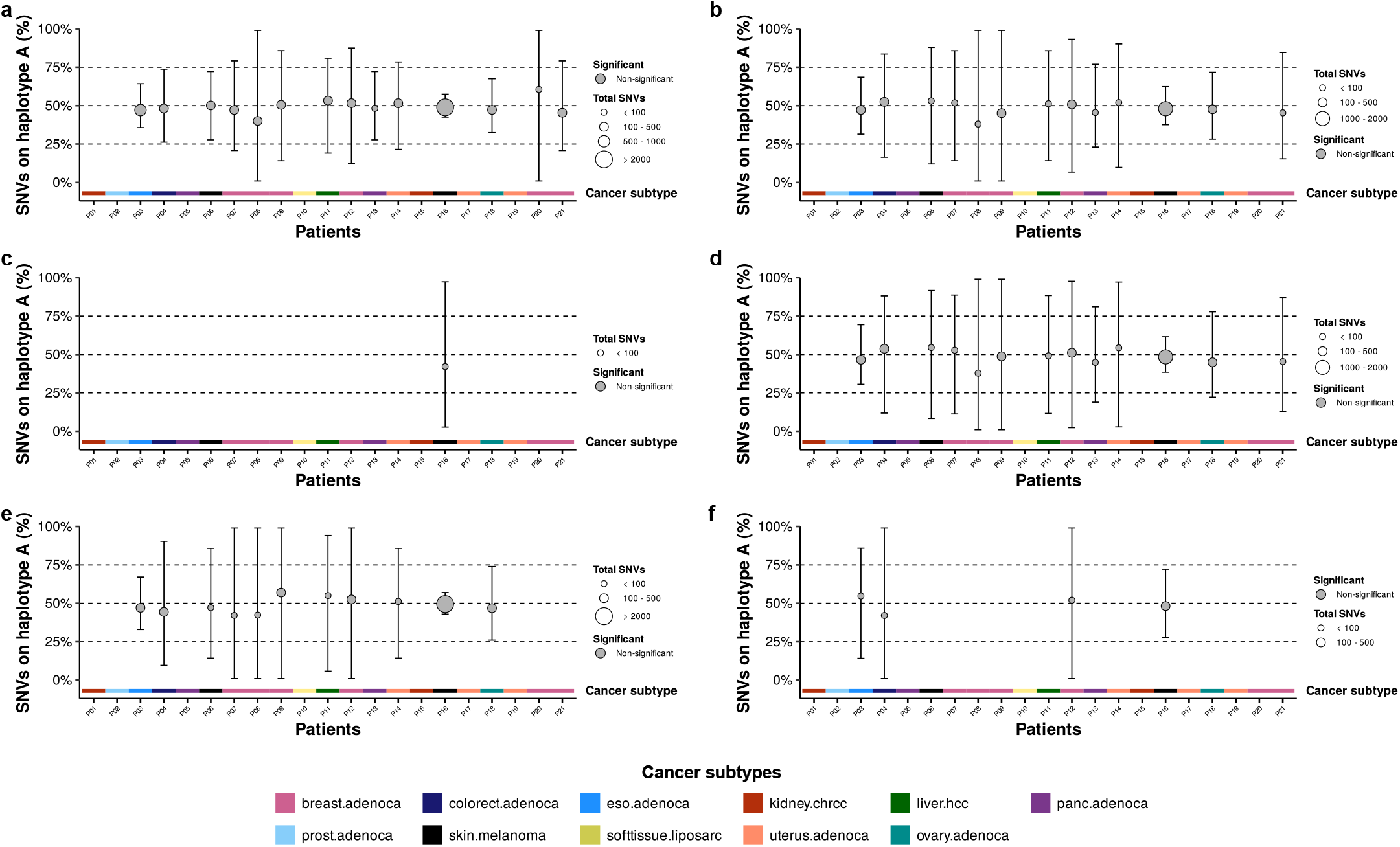
Very stable mutation accumulation across parental haplotypes in admixed PCAWG genomes in allelic imbalanced regions. Fraction of SNVs assigned to one haplotype across admixed patients with ≥50 SNVs. Each point represents a patient, sized by total SNVs;error bars show the maximum haplotype imbalance still consistent with each patient’s observed data (α = 0.05), accounting for phasing and local ancestry inference error. Dashed line at 50% indicates the expected equal distribution. Red points denote significant deviation from 50% after FDR correction (p < 0.05). **–** Genome-wide (autosomes) **(B)** Genes **(C)** Exons **(D)** Introns **(E)** Intergenic regions **(F)** C>TpG (SBS1) SNVs.

**Extended Data Figure S10.**
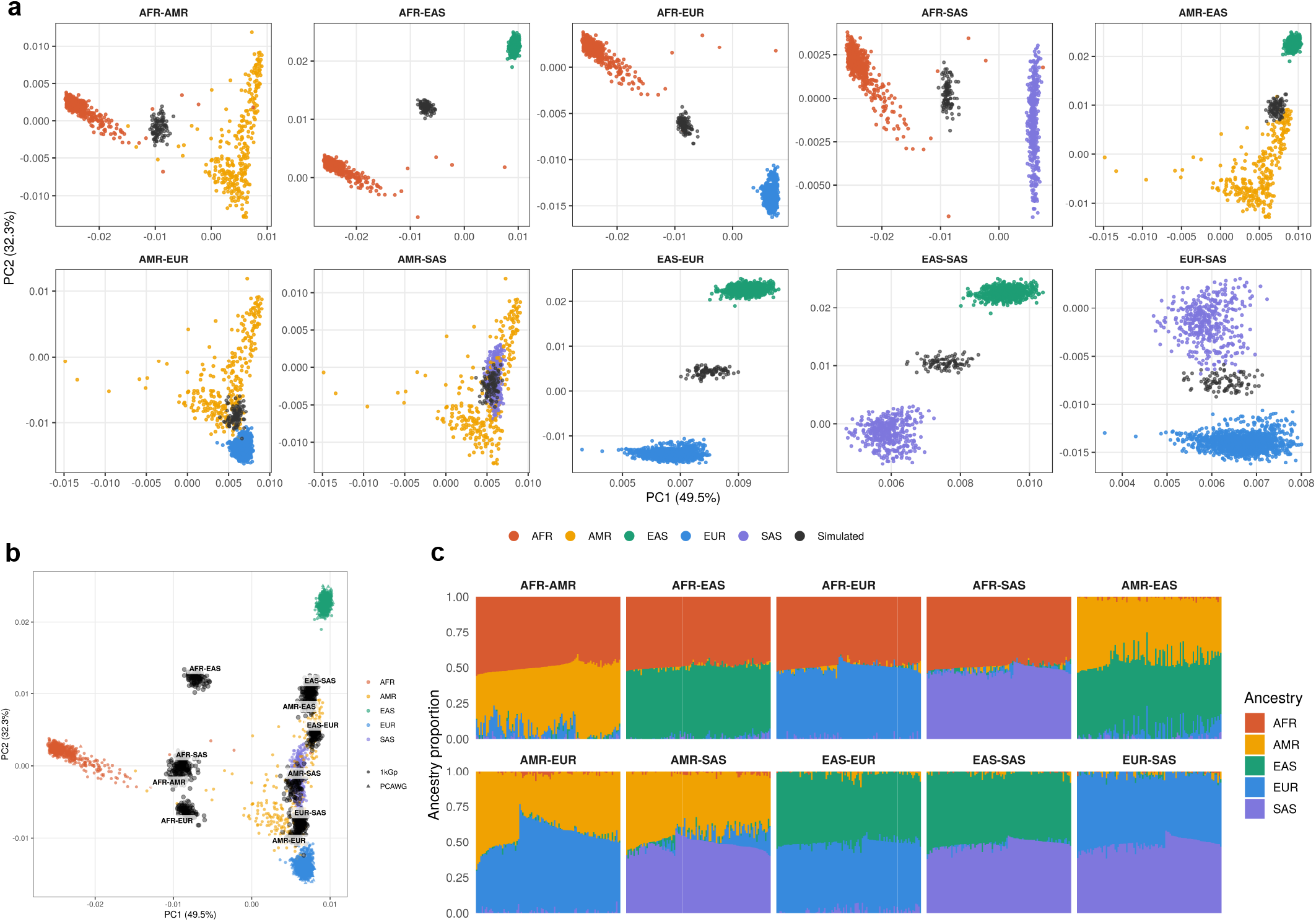
Global ancestry validation of simulated admixed individuals. **(A)** Principal component analysis on the 100,000 most discriminant SNPs, faceted by simulated ancestry combination; simulated individuals (grey) are shown against their corresponding pure 1kGp reference populations (colored). **(B)** Principal component analysis on the 100,000 most discriminant SNPs of 1kGp super-populations and simulated admixed individuals. 1kGp points are colored by self-reported ancestry; simulated individuals are shown in grey and labeled at the centroid of each simulated ancestry combination. Axes indicate variance explained by the first two PCs. **(C)** Global ancestry proportions of simulated individuals estimated by supervised ADMIXTURE, faceted by simulated ancestry combination. Each vertical bar represents one simulated individual; colors indicate the contribution of each super-population ancestry (AFR, AMR, EAS, EUR, SAS).

