## Supplementary Figures S1-20 for "Parent-of-origin phasing of somatic mutations shows equal mutation burden between parental genomes in human cancers"

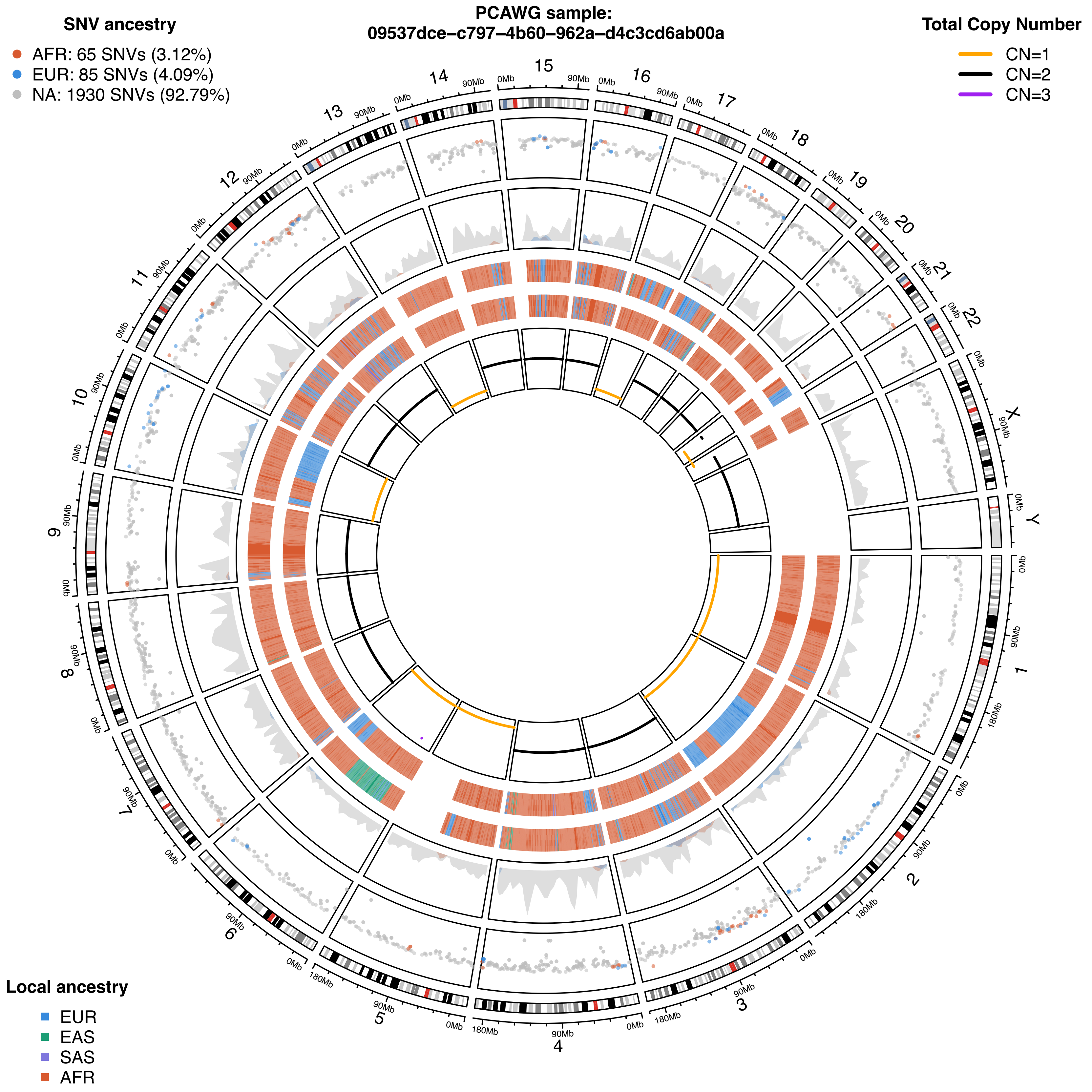
PCAWG sample:
09537dce−c797−4b60−962a−d4c3cd6ab00a
SNV ancestry
AFR: 65 SNVs (3.12%)
EUR: 85 SNVs (4.09%)
NA: 1930 SNVs (92.79%)
Total Copy Number
CN=1
CN=2
CN=3
Local ancestry
EUR
EAS
SAS
AFR
X
Y
0Mb
90Mb
180Mb

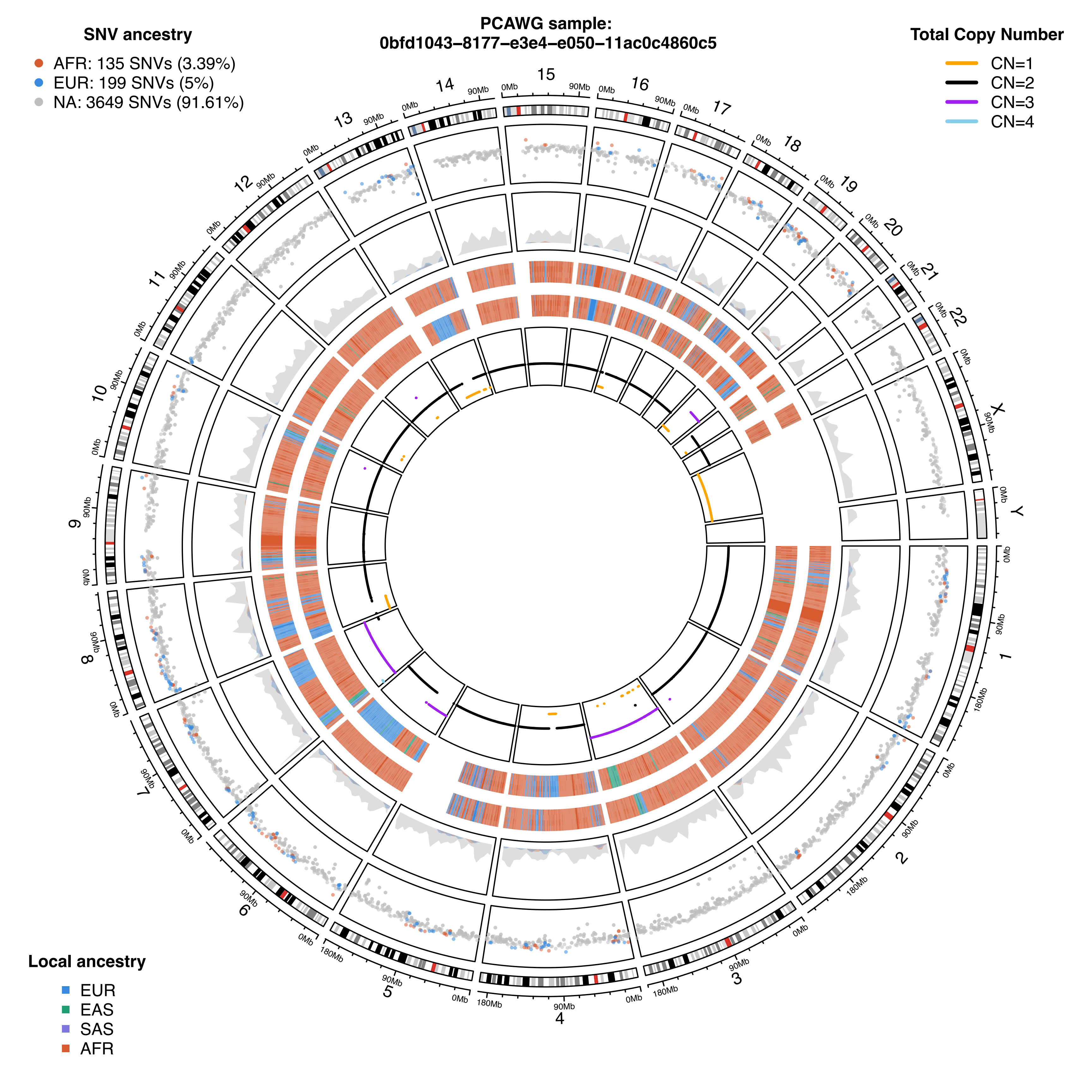

PCAWG sample:
0bfd1043–8177–e3e4–e050–11ac0c4860c5
SNV ancestry
AFR: 135 SNVs (3.39%)
EUR: 199 SNVs (5%)
NA: 3649 SNVs (91.61%)
Total Copy Number
CN=1
CN=2
CN=3
CN=4
Local ancestry
EUR
EAS
SAS
AFR

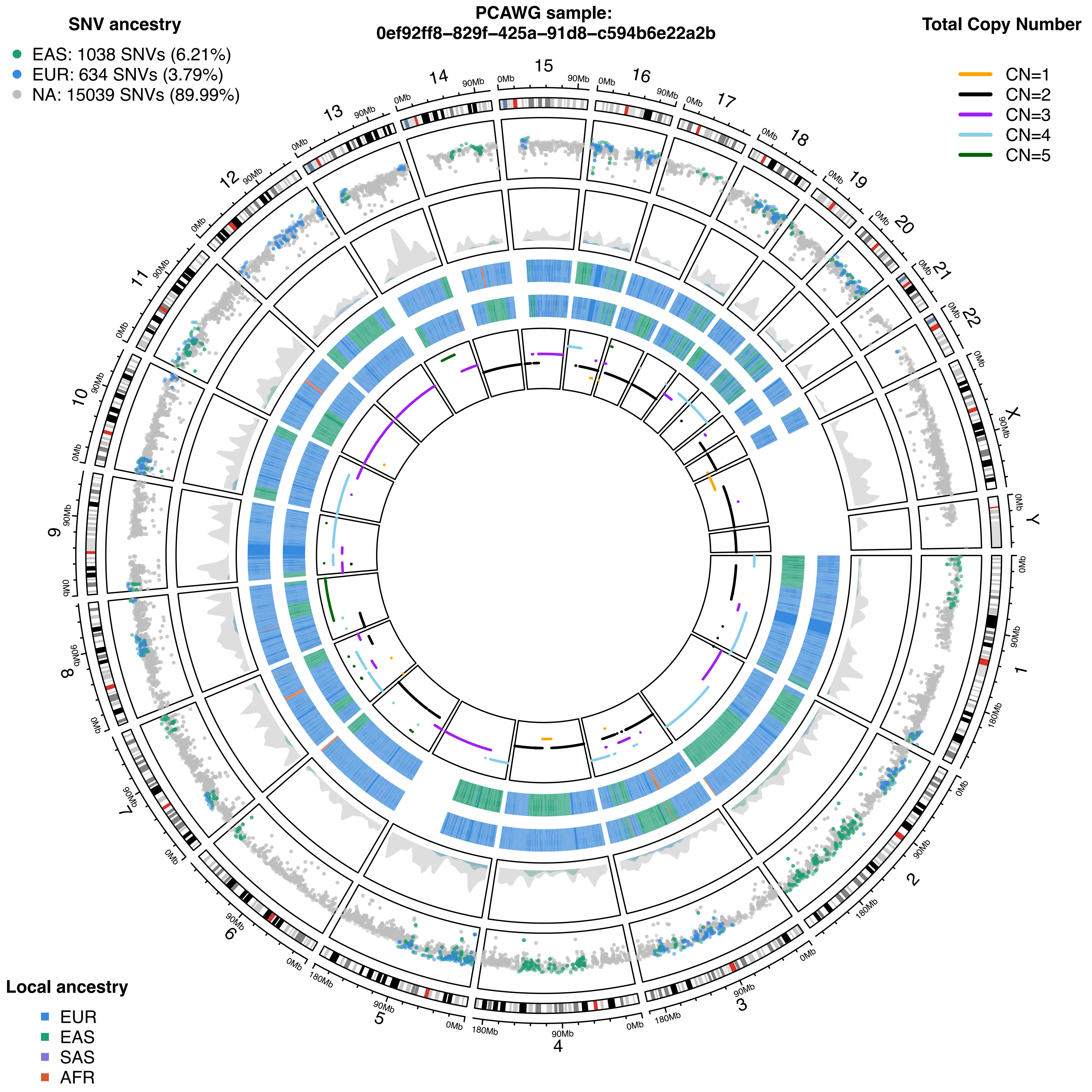

PCAWG sample:
0ef92ff8-829f-425a-91d8-c594b6e22a2b
SNV ancestry
EAS: 1038 SNVs (6.21%)
EUR: 634 SNVs (3.79%)
NA: 15039 SNVs (89.99%)
Total Copy Number
CN=1
CN=2
CN=3
CN=4
CN=5
Local ancestry
EUR
EAS
SAS
AFR
X
Y
0Mb
90Mb
180Mb

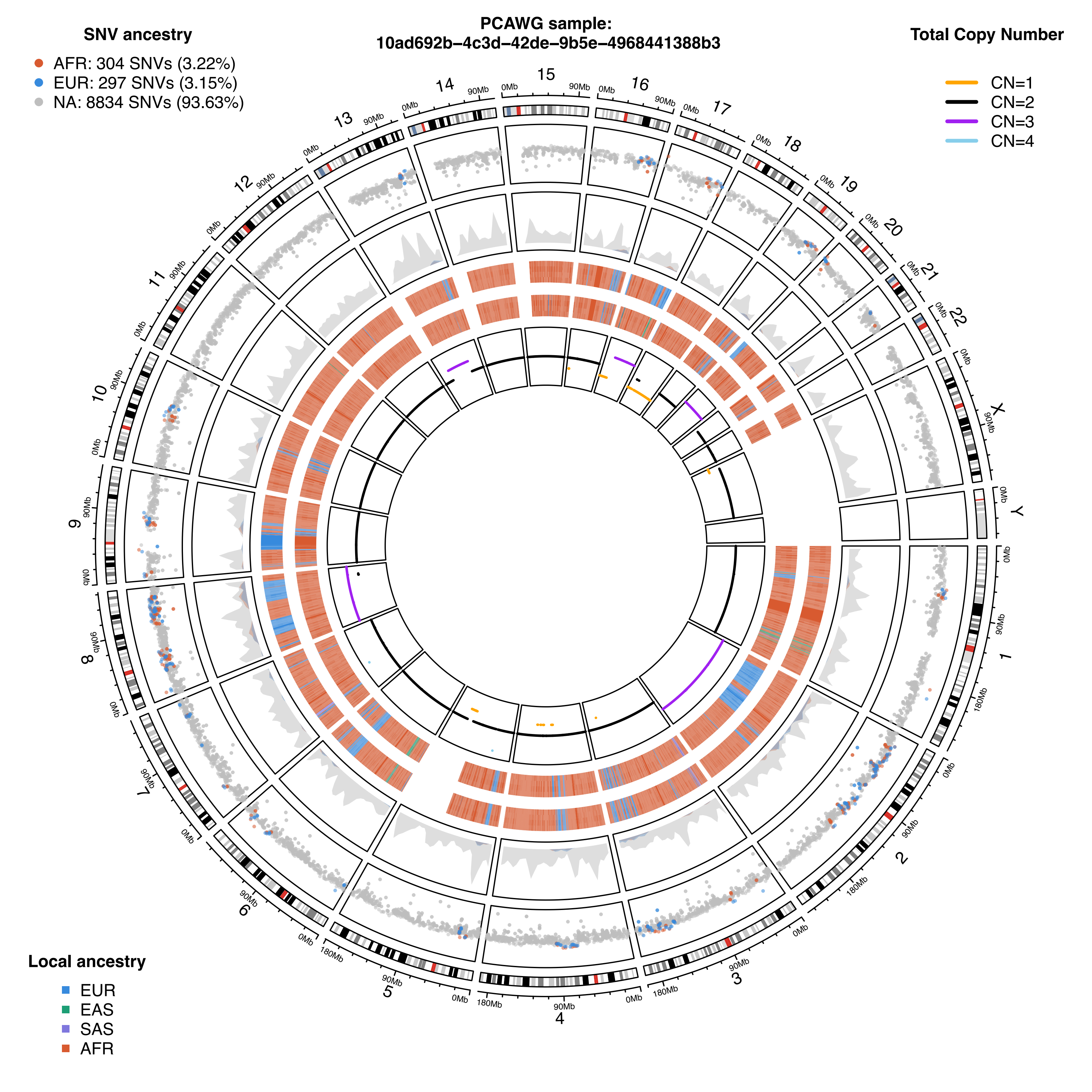

PCAWG sample:
10ad692b–4c3d–42de–9b5e–4968441388b3
SNV ancestry
AFR: 304 SNVs (3.22%)
EUR: 297 SNVs (3.15%)
NA: 8834 SNVs (93.63%)
Total Copy Number
CN=1
CN=2
CN=3
CN=4
Local ancestry
EUR
EAS
SAS
AFR

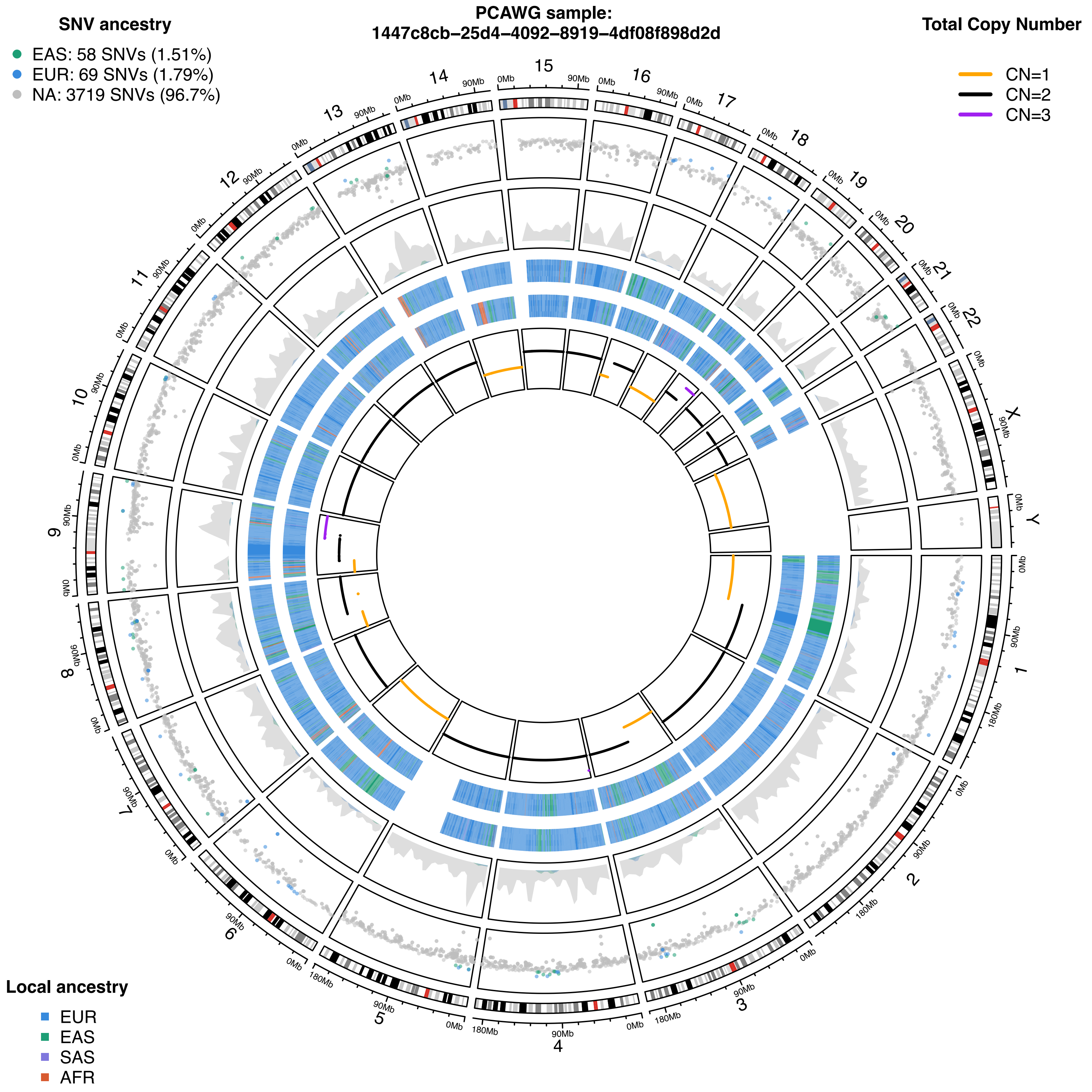

PCAWG sample:
1447c8cb-25d4-4092-8919-4df08f898d2d
SNV ancestry
EAS: 58 SNVs (1.51%)
EUR: 69 SNVs (1.79%)
NA: 3719 SNVs (96.7%)
Total Copy Number
CN=1
CN=2
CN=3
Local ancestry
EUR
EAS
SAS
AFR

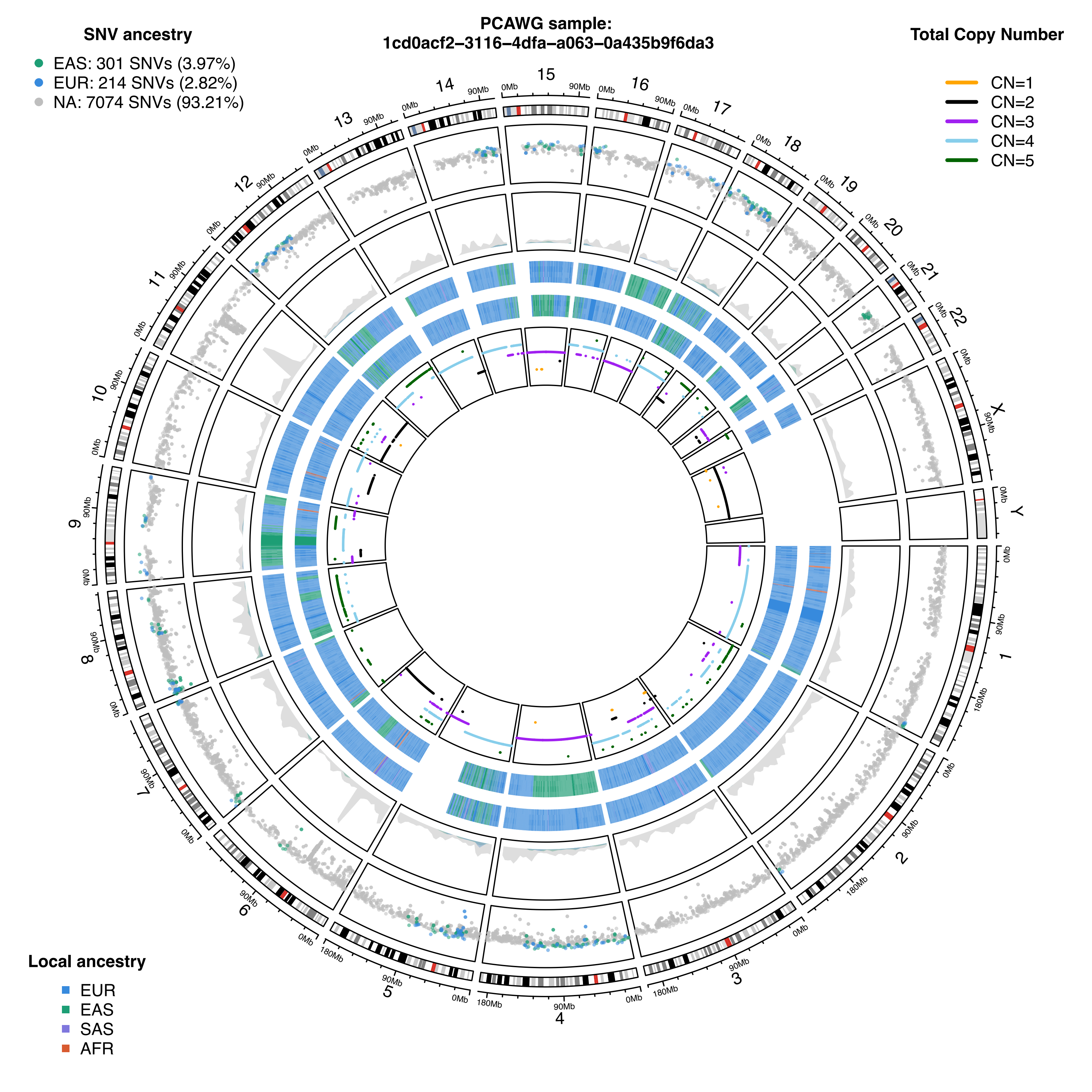
PCAWG sample:
1cd0acf2-3116-4dfa-a063-0a435b9f6da3
SNV ancestry
EAS: 301 SNVs (3.97%)
EUR: 214 SNVs (2.82%)
NA: 7074 SNVs (93.21%)
Total Copy Number
CN=1
CN=2
CN=3
CN=4
CN=5
Local ancestry
EUR
EAS
SAS
AFR

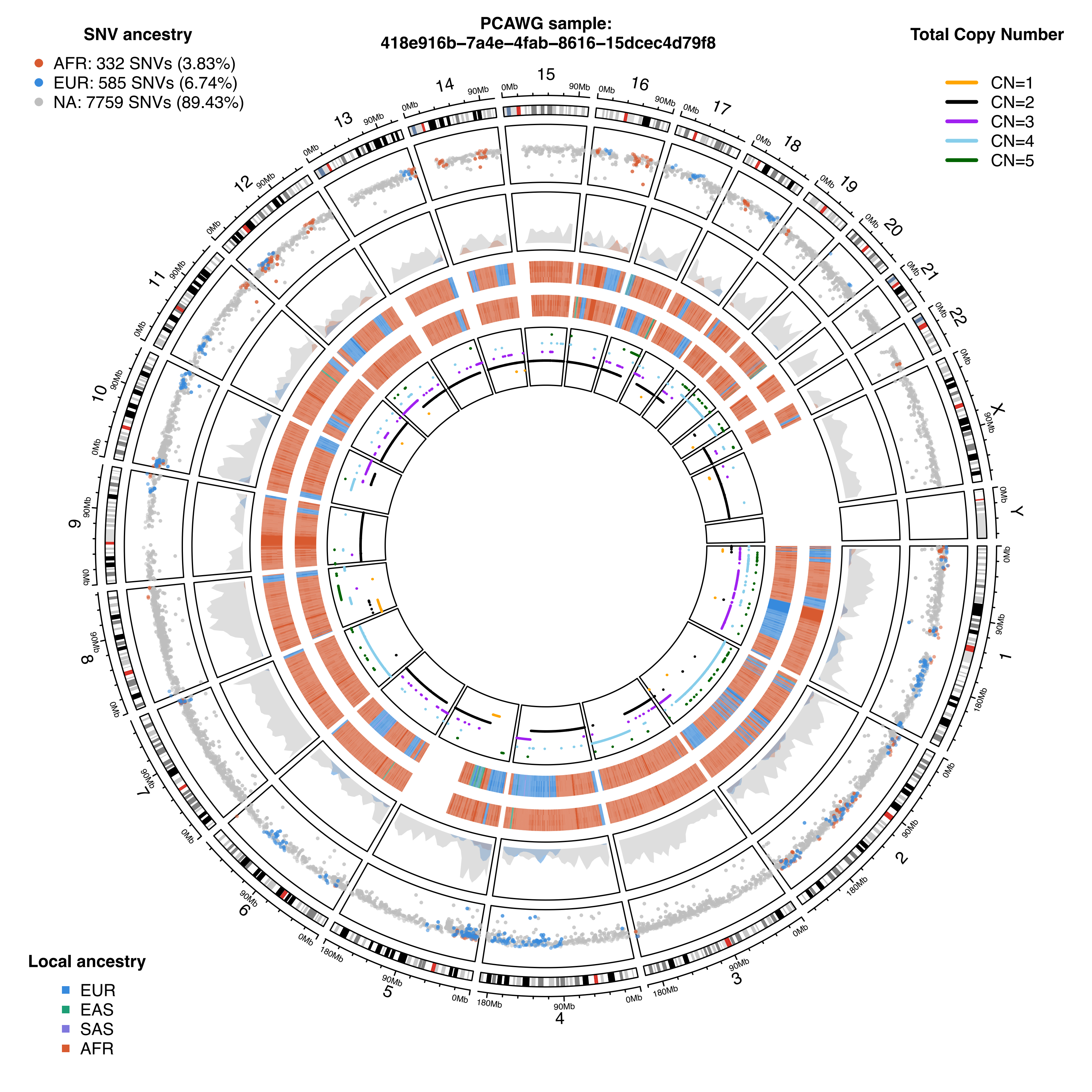

SNV ancestry
AFR: 332 SNVs (3.83%)
EUR: 585 SNVs (6.74%)
NA: 7759 SNVs (89.43%)
PCAWG sample:
418e916b−7a4e−4fab−8616−15dcec4d79f8
Total Copy Number
CN=1
CN=2
CN=3
CN=4
CN=5
Local ancestry
EUR
EAS
SAS
AFR

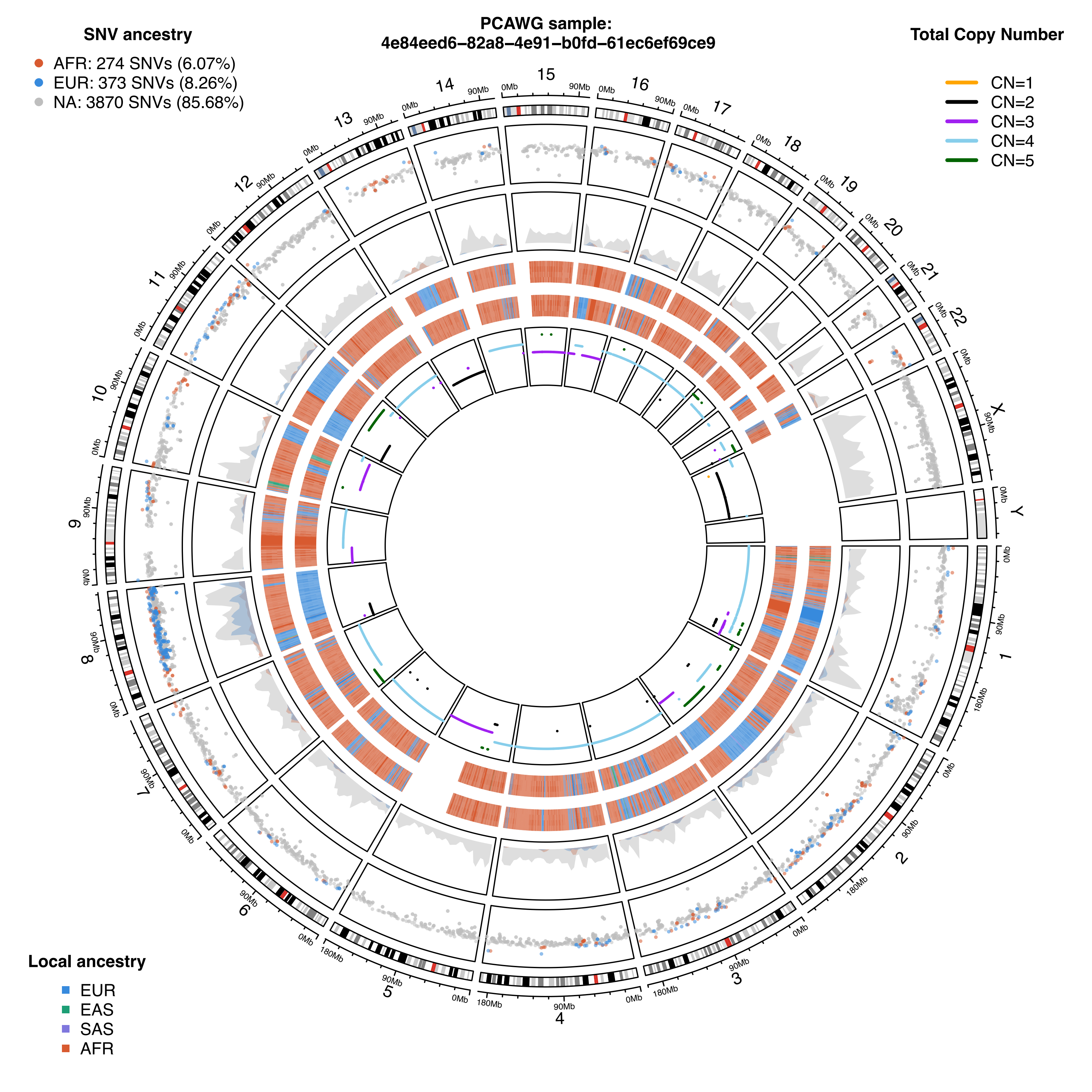
PCAWG sample:
4e84eed6–82a8–4e91–b0fd–61ec6ef69ce9
SNV ancestry
AFR: 274 SNVs (6.07%)
EUR: 373 SNVs (8.26%)
NA: 3870 SNVs (85.68%)
Total Copy Number
CN=1
CN=2
CN=3
CN=4
CN=5
Local ancestry
EUR
EAS
SAS
AFR

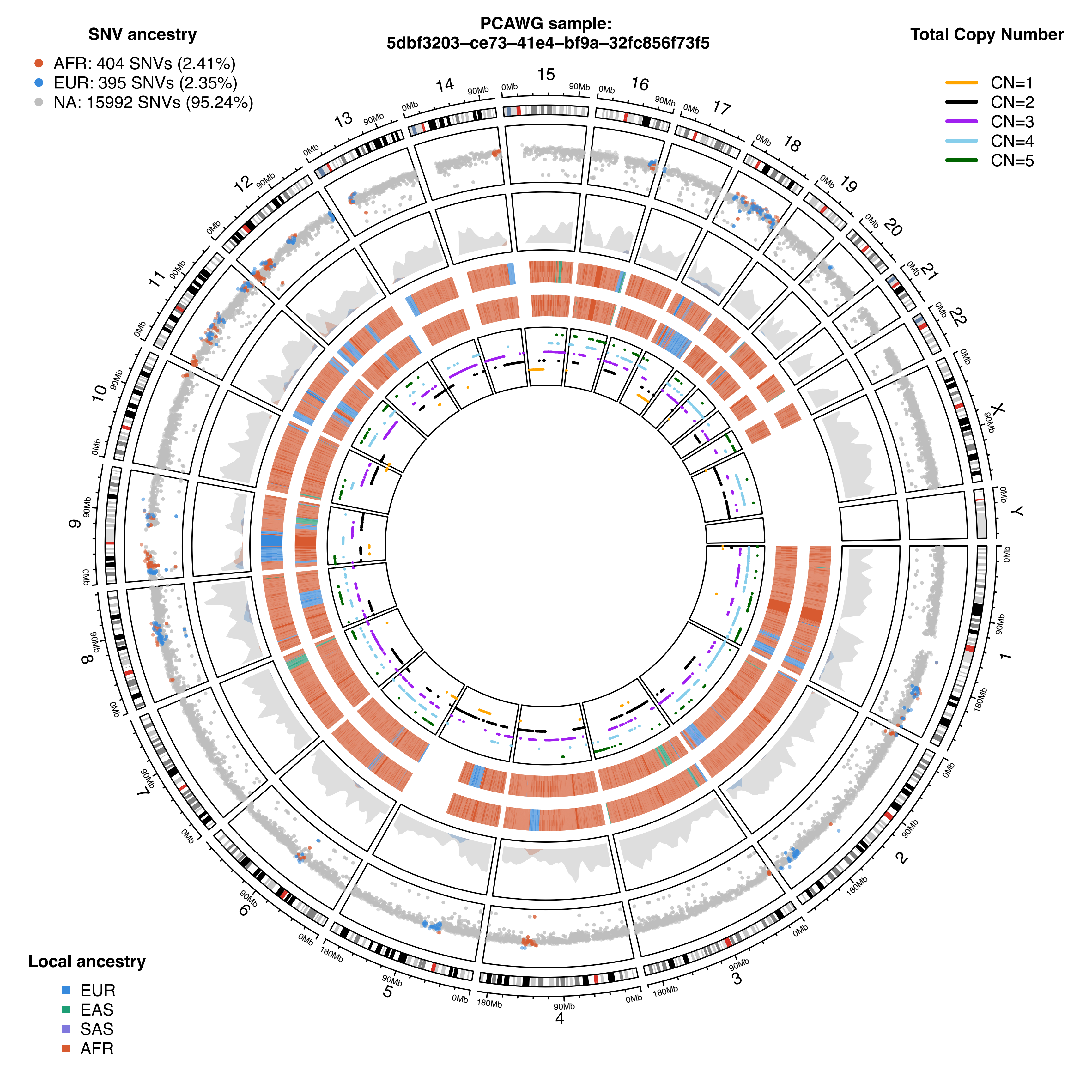

PCAWG sample:
5dbf3203-ce73-41e4-bf9a-32fc856f73f5
SNV ancestry
AFR: 404 SNVs (2.41%)
EUR: 395 SNVs (2.35%)
NA: 15992 SNVs (95.24%)
Total Copy Number
CN=1
CN=2
CN=3
CN=4
CN=5
Local ancestry
EUR
EAS
SAS
AFR

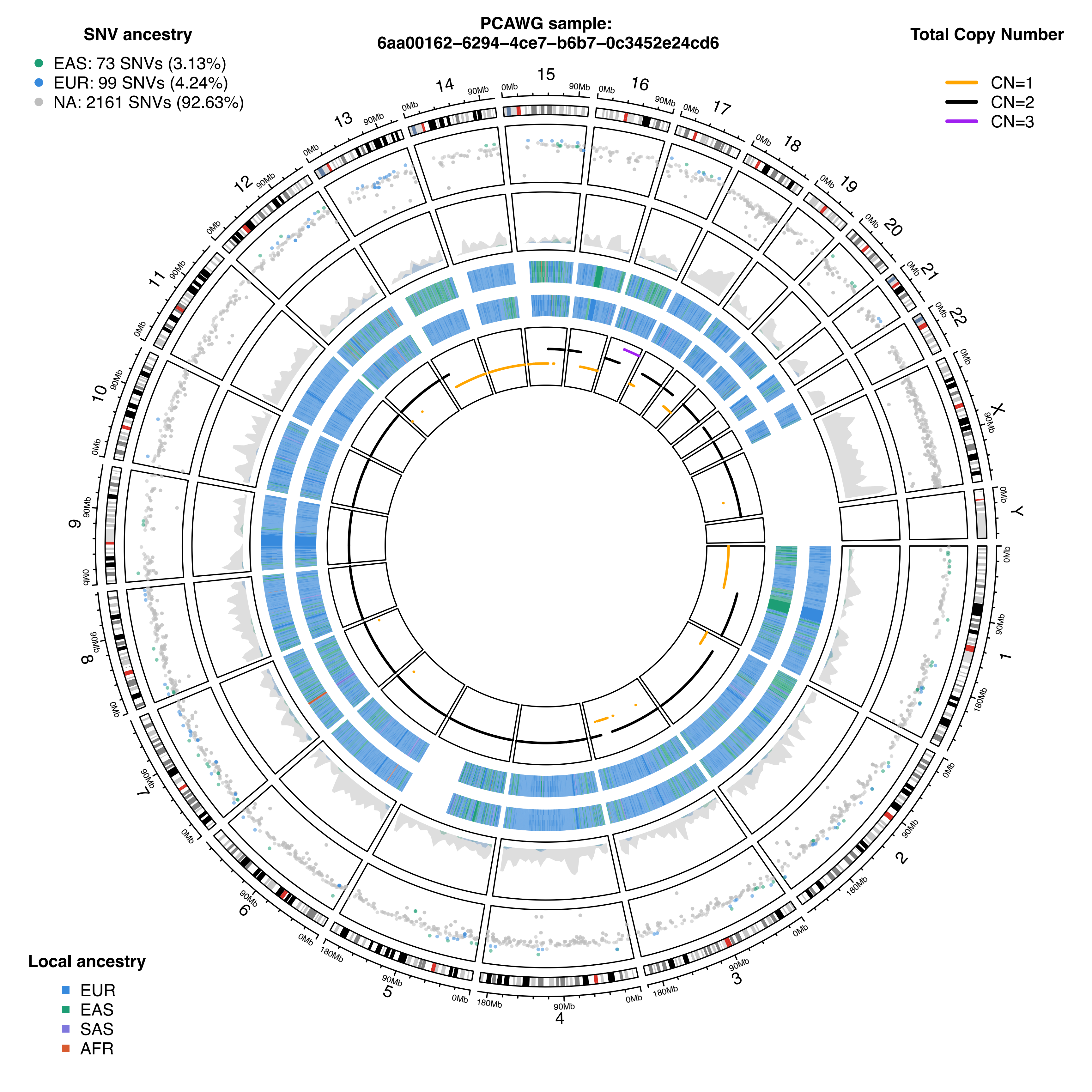
PCAWG sample:
6aa00162–6294–4ce7–b6b7–0c3452e24cd6
SNV ancestry
EAS: 73 SNVs (3.13%)
EUR: 99 SNVs (4.24%)
NA: 2161 SNVs (92.63%)
Total Copy Number
CN=1
CN=2
CN=3
Local ancestry
EUR
EAS
SAS
AFR
X
Y
0Mb
90Mb
180Mb

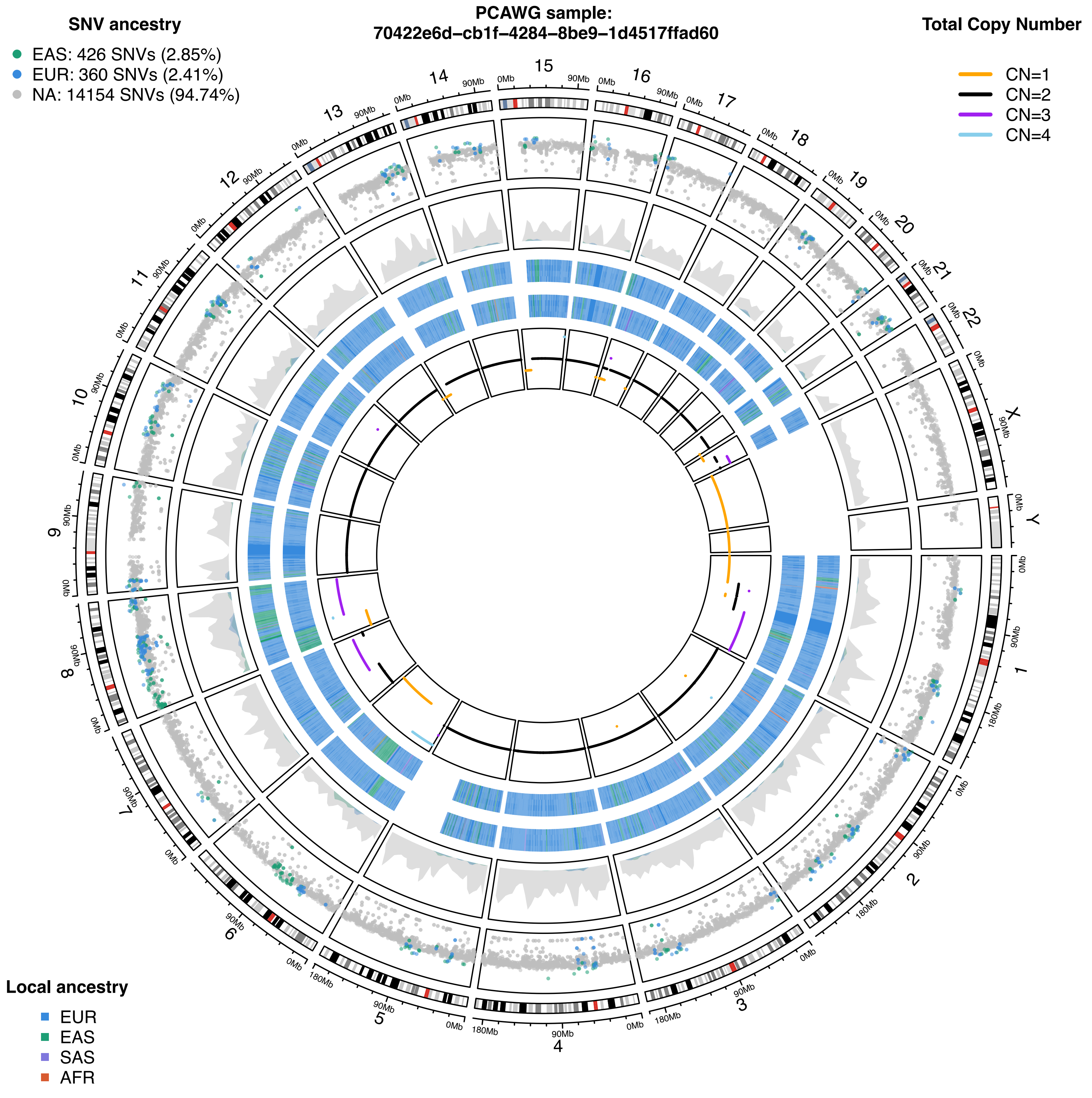

PCAWG sample:
70422e6d–cb1f–4284–8be9–1d4517ffad60
SNV ancestry
EAS: 426 SNVs (2.85%)
EUR: 360 SNVs (2.41%)
NA: 14154 SNVs (94.74%)
Total Copy Number
CN=1
CN=2
CN=3
CN=4
Local ancestry
EUR
EAS
SAS
AFR
X
Y
0Mb
90Mb
180Mb

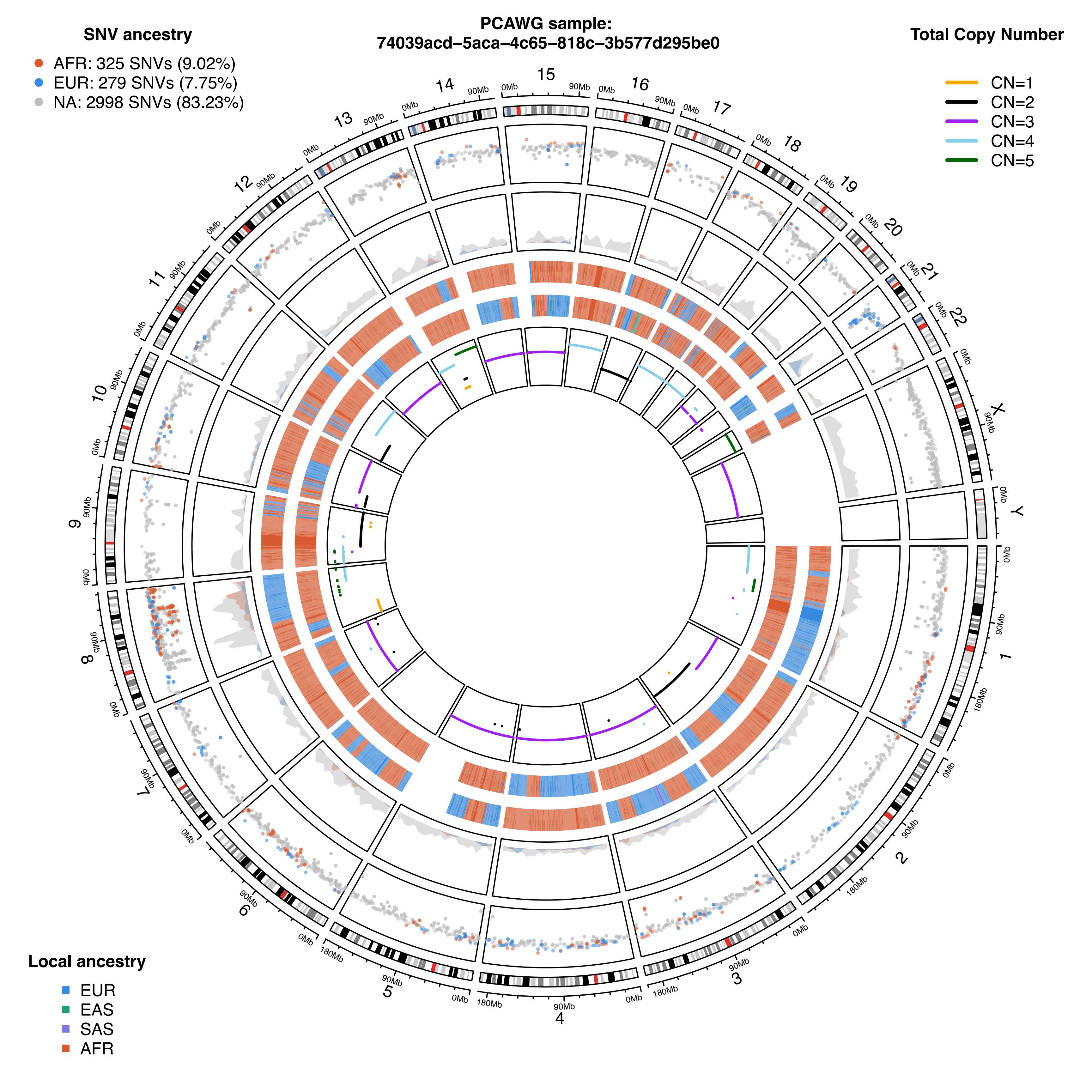

PCAWG sample:
74039acd–5aca–4c65–818c–3b577d295be0
SNV ancestry
AFR: 325 SNVs (9.02%)
EUR: 279 SNVs (7.75%)
NA: 2998 SNVs (83.23%)
Total Copy Number
CN=1
CN=2
CN=3
CN=4
CN=5
Local ancestry
EUR
EAS
SAS
AFR

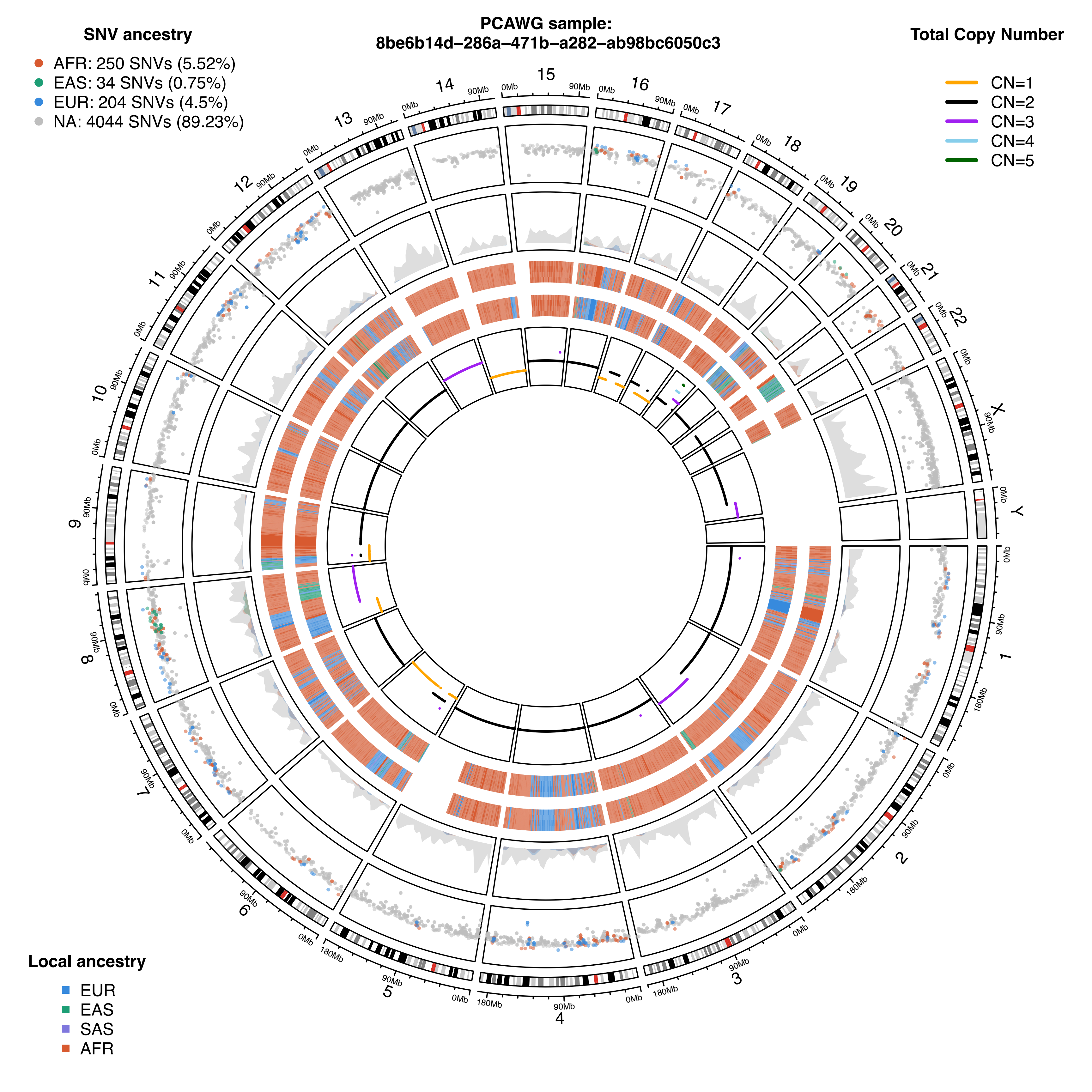
PCAWG sample:
8be6b14d-286a-471b-a282-ab98bc6050c3
SNV ancestry
AFR: 250 SNVs (5.52%)
EAS: 34 SNVs (0.75%)
EUR: 204 SNVs (4.5%)
NA: 4044 SNVs (89.23%)
Total Copy Number
CN=1
CN=2
CN=3
CN=4
CN=5
Local ancestry
EUR
EAS
SAS
AFR

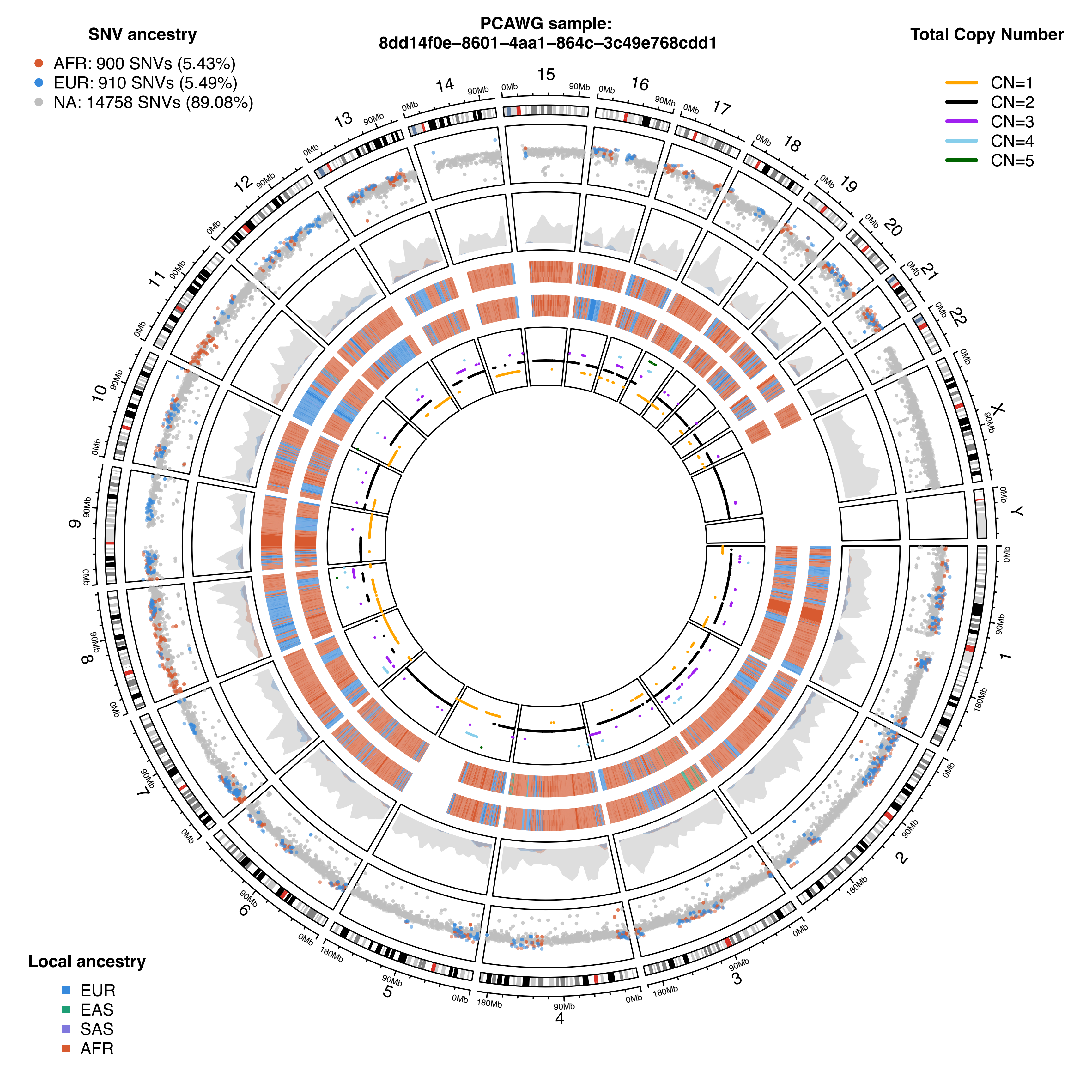

SNV ancestry
AFR: 900 SNVs (5.43%)
EUR: 910 SNVs (5.49%)
NA: 14758 SNVs (89.08%)
PCAWG sample:
8dd14f0e-8601-4aa1-864c-3c49e768cdd1
Total Copy Number
CN=1
CN=2
CN=3
CN=4
CN=5
Local ancestry
EUR
EAS
SAS
AFR

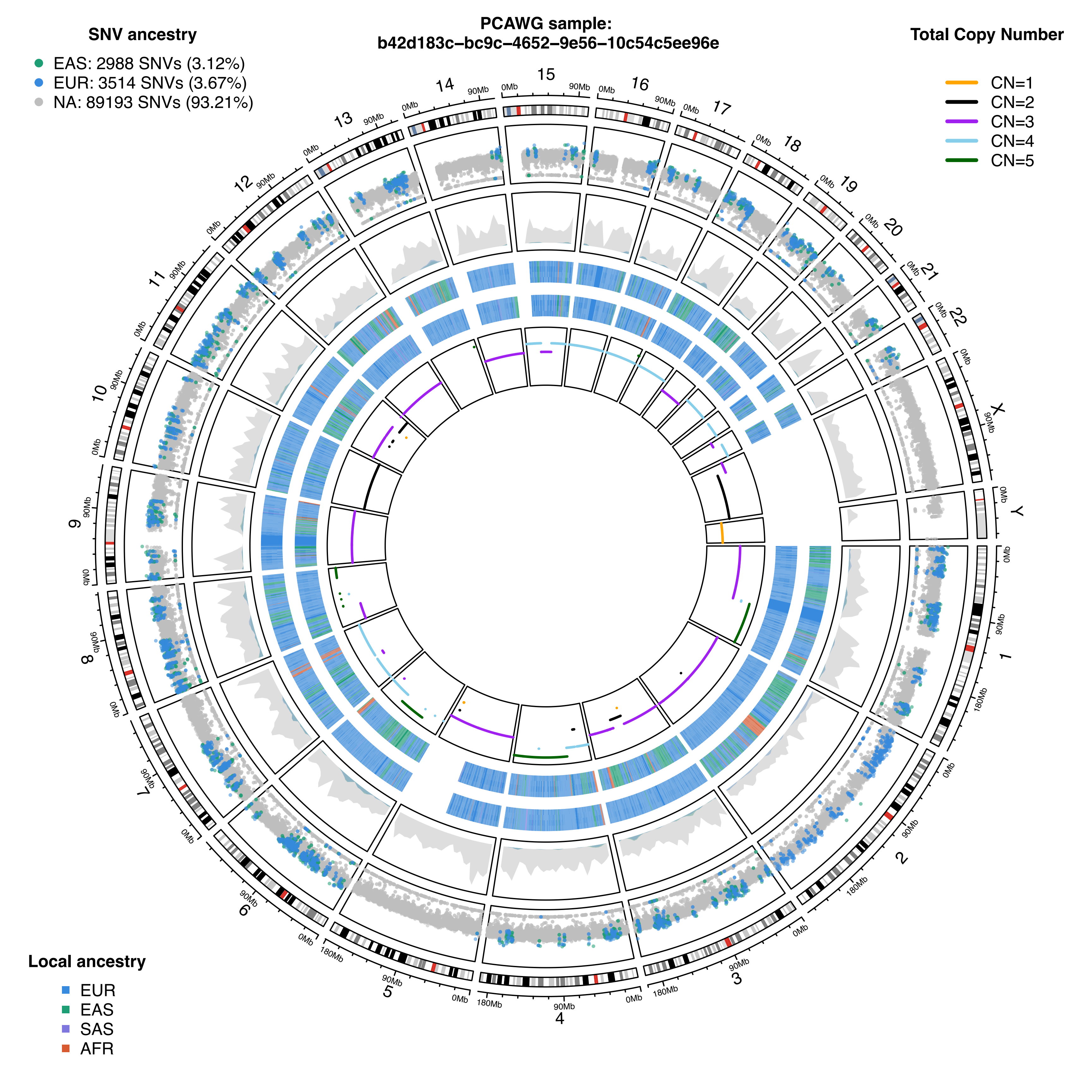
PCAWG sample:
b42d183c-bc9c-4652-9e56-10c54c5ee96e
SNV ancestry
EAS: 2988 SNVs (3.12%)
EUR: 3514 SNVs (3.67%)
NA: 89193 SNVs (93.21%)
Total Copy Number
CN=1
CN=2
CN=3
CN=4
CN=5
Local ancestry
EUR
EAS
SAS
AFR

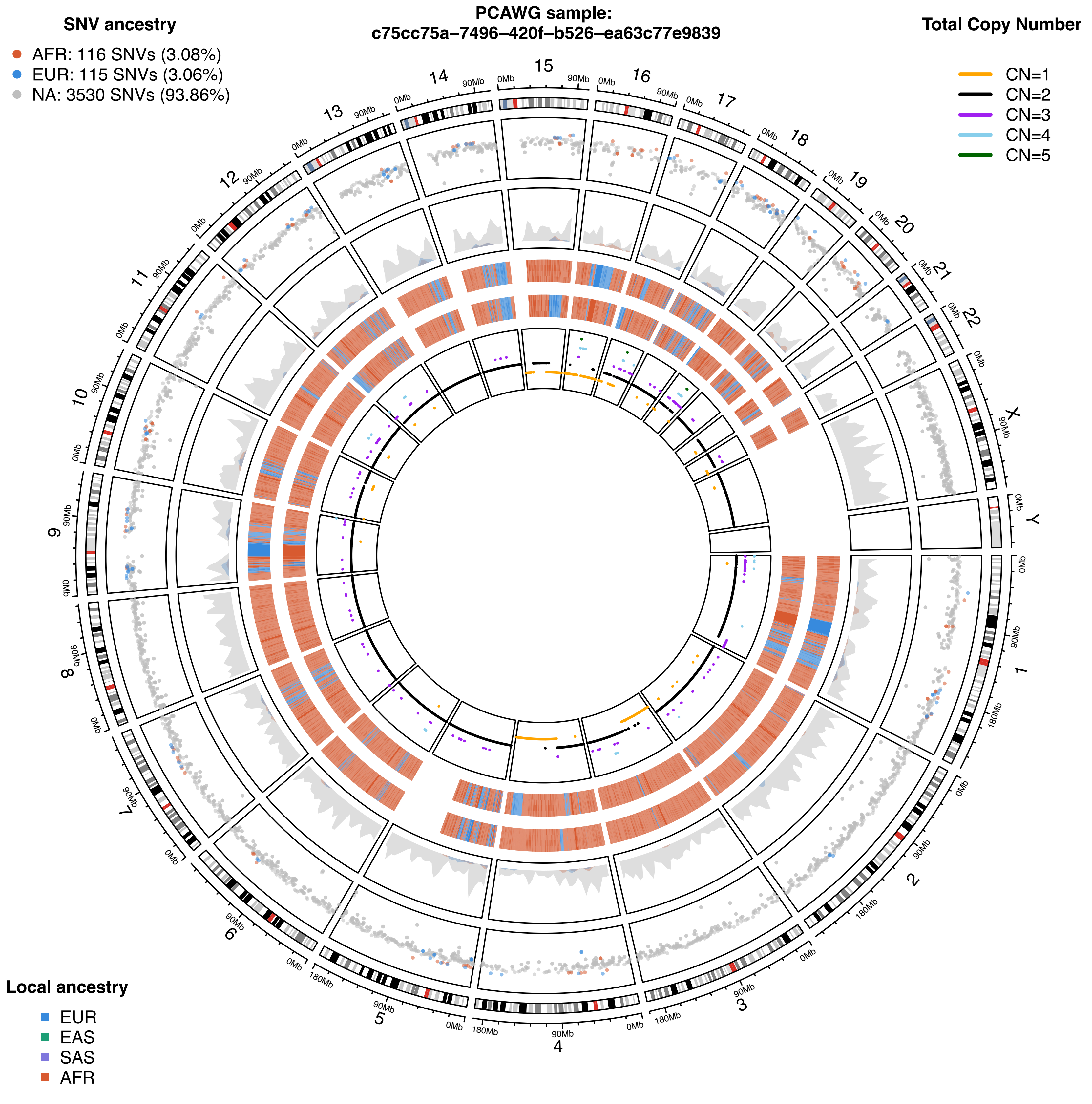
PCAWG sample:
c75cc75a-7496-420f-b526-ea63c77e9839
SNV ancestry
AFR: 116 SNVs (3.08%)
EUR: 115 SNVs (3.06%)
NA: 3530 SNVs (93.86%)
Total Copy Number
CN=1
CN=2
CN=3
CN=4
CN=5
Local ancestry
EUR
EAS
SAS
AFR

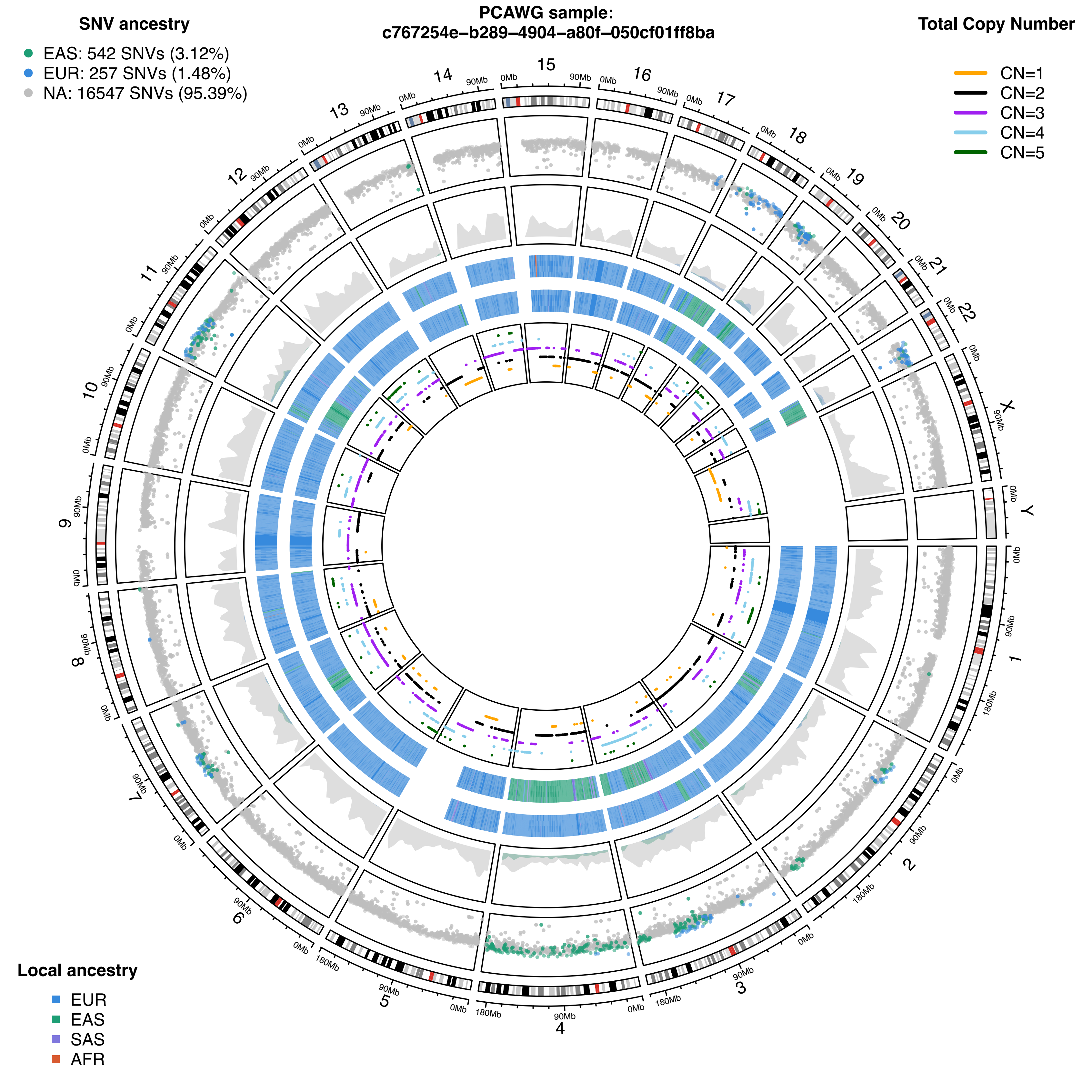
PCAWG sample:
c767254e–b289–4904–a80f–050cf01ff8ba
SNV ancestry
EAS: 542 SNVs (3.12%)
EUR: 257 SNVs (1.48%)
NA: 16547 SNVs (95.39%)
Total Copy Number
CN=1
CN=2
CN=3
CN=4
CN=5
Local ancestry
EUR
EAS
SAS
AFR
X
Y
0Mb
90Mb
180Mb

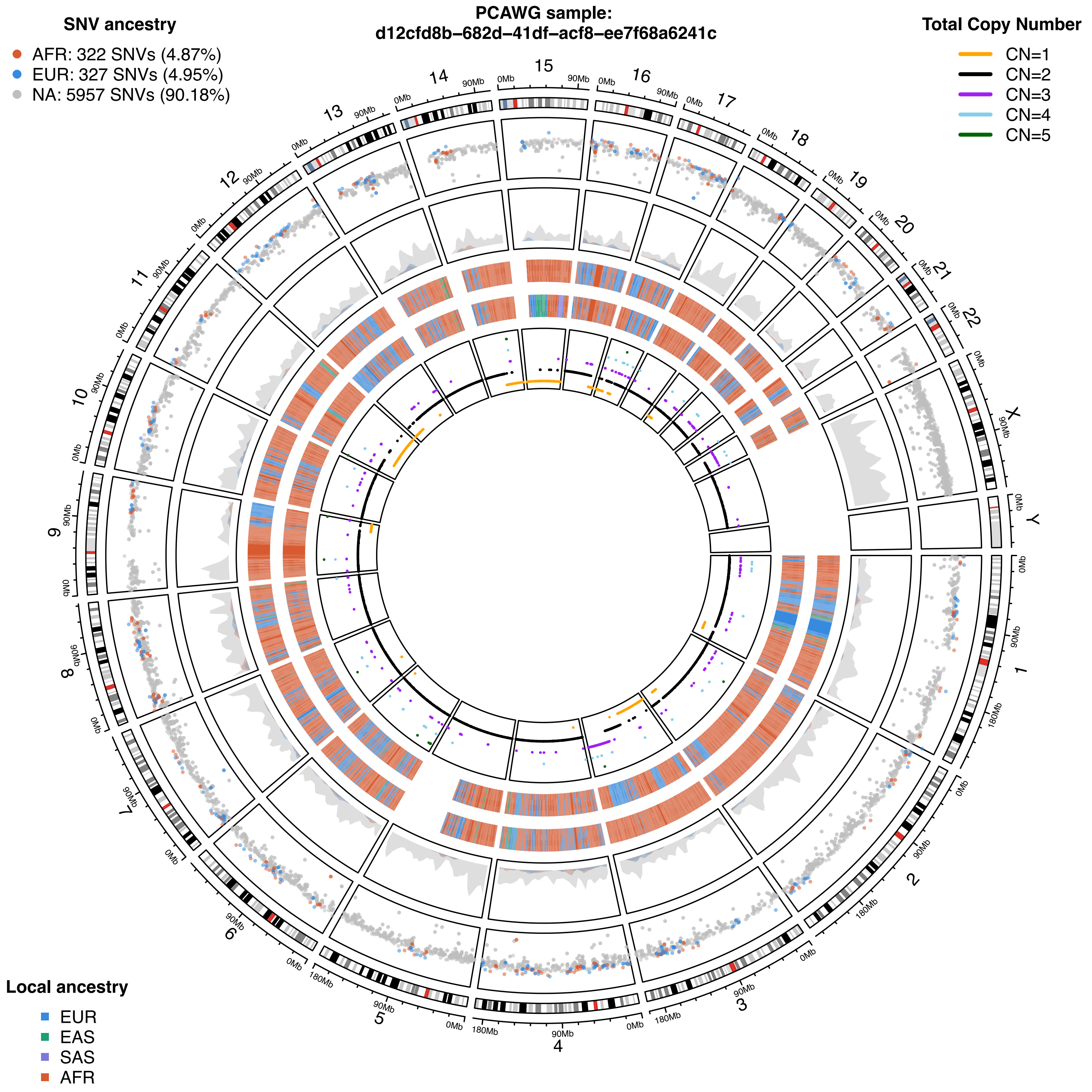
PCAWG sample:
d12cfd8b-682d-41df-acf8-ee7f68a6241c
SNV ancestry
AFR: 322 SNVs (4.87%)
EUR: 327 SNVs (4.95%)
NA: 5957 SNVs (90.18%)
Total Copy Number
CN=1
CN=2
CN=3
CN=4
CN=5
Local ancestry
EUR
EAS
SAS
AFR
X
Y
0Mb
90Mb
180Mb

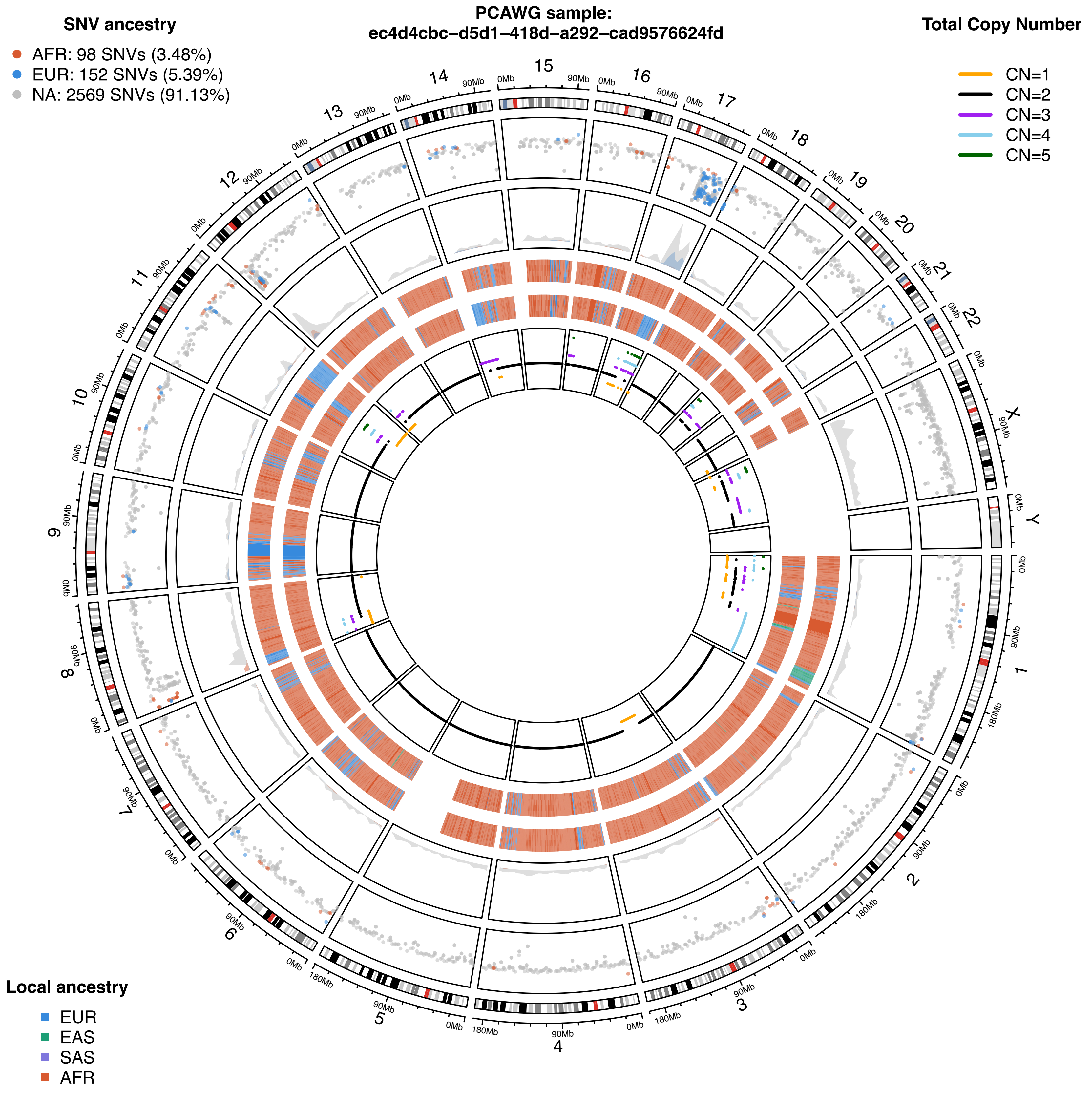
PCAWG sample:
ec4d4cbc-d5d1-418d-a292-cad9576624fd
SNV ancestry
AFR: 98 SNVs (3.48%)
EUR: 152 SNVs (5.39%)
NA: 2569 SNVs (91.13%)
Total Copy Number
CN=1
CN=2
CN=3
CN=4
CN=5
Local ancestry
EUR
EAS
SAS
AFR

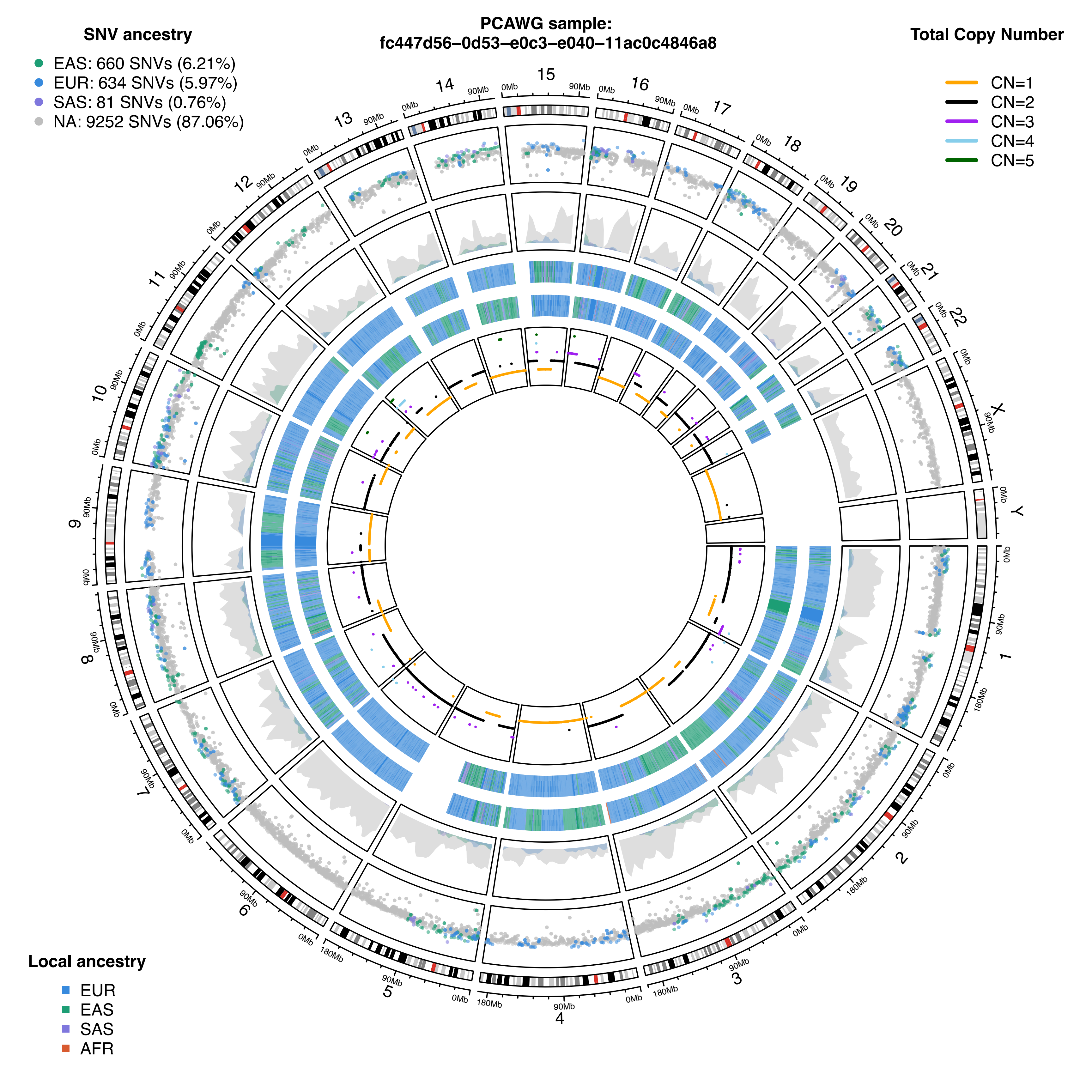
PCAWG sample:
fc447d56–0d53–e0c3–e040–11ac0c4846a8
SNV ancestry
EAS: 660 SNVs (6.21%)
EUR: 634 SNVs (5.97%)
SAS: 81 SNVs (0.76%)
NA: 9252 SNVs (87.06%)
Total Copy Number
CN=1
CN=2
CN=3
CN=4
CN=5
Local ancestry
EUR
EAS
SAS
AFR
X
Y
0Mb
90Mb
180Mb
