## Supplementary Info-Table legends for "Parent-of-origin phasing of somatic mutations shows equal mutation burden between parental genomes in human cancers"

**Supplementary Table 1. Overview of the 21 informative admixed PCAWG patients.** SNV counts at each pipeline stage (phased, assignable, assigned to haplotype A/B in balanced and imbalanced regions), haplotype ancestries, sex, and resolved maternal/paternal identity where available.

**Supplementary Table 2. Validation of the copy-number correction for SNV quantification.**
Slope of mutation count versus major copy number, before and after correction, estimated from simulations and from imbalanced PCAWG segments (major copy number 2–5).

**Supplementary Table 3. Retrospective compatibility bounds on parental haplotype imbalance.**
Per-patient local ancestry accuracy, copy-number-corrected haplotype A/B mutation counts, binomial test p-value, and the maximum haplotype imbalance compatible with the data at $\alpha = 0.05$.

**Supplementary information. Local ancestry profile of admixed informative patients inferred with Gnomix.**

Tracks (inner to outer): total copy number, haplotype-level local ancestry, ancestry-specific SNV density, and individual SNVs by ancestry
